# Conventional dendritic cells type I with an enhanced type-I-IFN signaling underpin anti-tumor immune responses in brain metastases

**DOI:** 10.64898/2026.07.08.737181

**Authors:** Fiona James, Alena Revalova, Christopher Fife, Jennifer Williams, Dario Guglietta, Zarnaz Hadi, Elton J. R. Vasconcelos, Ashley Sunderland, Grace Mallett, Nicola Ingram, Tsuneyasu Kaisho, William J. Brackenbury, Merin Lawrence, David R. Westhead, Andrew S. MacDonald, Mihaela Lorger

## Abstract

Brain metastases are associated with poor prognosis. A better understanding of anti-tumor immune responses in the context of immune specialized microenvironment of the brain is required to develop improved therapeutic strategies for this disease. We demonstrate that the conventional dendritic cells type 1 (cDC1) gene signature positively correlates with a prolonged BrM-dependent survival in melanoma and breast cancer patients. Furthermore, intracranial anti-tumor immune responses in preclinical BrM models consistently rely on cDC1s for tumor growth control, BrM-dependent survival and maintenance of the intra-tumoral CD8+ T cell pool, in contrast to variable, cancer type-dependent cDC1 roles in extracranial tumors. This is underpinned by tumor site-specific cDC1 molecular profiles with distinct Toll like receptor repertoires, upregulation of co-stimulatory molecules and IL-12, and enhanced type-I-IFN signaling in intracranial cDC1s, with the latter driving increased cDC1 activation. cDC1s also promote the conversion of progenitor exhausted CD8+ T cells to transient effectors, which is further enhanced by immune checkpoint blockade therapy. These findings pinpoint cDC1s as a major cell population of interest in the development of future immunotherapies for BrM.

**Significance statement:** Characteristics of immune cells and tumor microenvironment in brain metastases (BrM) are distinct to extracranial tumor sites and characterized by special immunological barriers and suppressive stroma. Conventional dendritic cells type 1 (cDC1s), which are key regulators of anti-tumor immunity, remain largely unexplored in this context. This study deepens our understanding of how anti-tumor immune responses differ between BrM and extracranial tumors by revealing BrM-specific phenotype and function of cDC1s, with the latter being critical for anti-tumor immunity in BrM even in cancer models where extracranial cDC1s demonstrate pro-tumorigenic function, highlighting cDC1s as universal regulators of intracranial anti-tumor immunity. Our study suggests that tumor site-specific augmentation of cDC1s should be pursued to enhance the efficacy of immunotherapies in BrM.

## Introduction

Brain metastases (BrM) develop in ∼30% of patients with metastatic cancer and predominantly originate from melanoma, breast and lung cancer. While treatment options for BrM vary depending on the cancer type, they are generally suboptimal and lag behind therapies for extracranial disease (1). Immune checkpoint blockade (ICB) is becoming increasingly important in treatment of BrM, with a combined PD-1/CTLA-4 blockade demonstrating a superior activity over monotherapies and reaching intracranial response rates of 45-55% in melanoma patients with asymptomatic BrM (2, 3). While exclusion of patients with breast cancer BrM from clinical trials (4) limits the data available, the activity in breast cancer appears lower.

The brain, in contrast to extracranial sites, is an immune-specialized environment with limited anti-tumor immune responses. A better understanding of anti-tumor immunity in the brain is required to support the development of improved strategies against BrM. Dendritic cells are the major antigen presenting cell population and a key component of the immune system, with CD103+Xcr1+ conventional dendritic cells type 1 (cDC1s) prominently involved in anti-tumor immune responses at extracranial sites, including in the context of ICB (5–7). cDC1s transport antigens from tumors to the tumor draining lymph nodes (TDLNs) and are critical for the priming of CD8+ T cell responses. The initial activation of CD8+ T cells in TDLNs gives rise to progenitor exhausted T cells (Tpex) that then migrate to tumors (8), while a reservoir of tumor antigen-specific Tpex is simultaneously maintained within TDLNs with help of cDC1s (9). Tpex conversion to transient effectors (Teff) with cytotoxic function and subsequently to terminally exhausted T cells (Tex) occur within tumors and require additional co-stimulation from antigen presenting cells (8, 10). In contrast to extracranial tumors, the role of cDC1s in the context of immune-specialized microenvironment in BrM remains largely unexplored. Addressing this knowledge gap, our study demonstrates a higher dependence of intracranial as compared to extracranial anti-tumor immunity on cDC1s and identifies potential underlying mechanisms, providing a rationale for focusing on cDC1s for improved immunotherapeutic strategies for BrM.

## Results

### cDC1 population consistently restricts tumor progression in the brain despite variable role in extracranial tumors

Analysis of publicly available mRNAseq data of human BrM (11, 12) revealed that high expression of cDC1 gene signature (consisting of XCR1 and BATF3) in BrM significantly correlates with a longer OS from craniotomy in melanoma, and with a better BrM-specific survival in breast cancer (**Fig. 1A**), suggesting a positive role for cDC1s in the control of BrM in patients. To investigate this further, we used our previously established two-site BrM model in which mice are bearing extracranial tumors in addition to tumors in the brain (13) (**Fig. 1B**). This mimics the presence of extracranial disease in patients and is required to reproduce the clinically observed intracranial ICB efficacy (2, 3, 13). The presence of cDC1s and cDC2s in intracranial B16-OVA tumors was confirmed by flow cytometry (**Fig. 1C; Fig. S1A-D**) and CD103+MHCII+ DCs could be also detected by immunofluorescence (**Fig. S1E**). The functional role of cDC1s was then assessed in Batf3-/-mice, which lack cDC1s due to the deletion of the transcription factor Batf3 essential for cDC1 development (14) without affecting cDC2 abundance (**Fig. S1F, G**). Intracranial B16-OVA melanoma tumor burden was significantly higher in Batf3-/- than WT mice, while the extracranial tumor burden was comparable (**Fig. 1D, E**). In line with this, the intracranial tumor-dependent survival in the two-site model was significantly shorter in the absence of cDC1s, while the extracranial tumor-dependent survival, quantified in mice bearing extracranial tumors only, was unaffected (**Fig. 1F**). In the RET melanoma two-site model, both intracranial and extracranial tumor growth were enhanced in the absence of cDC1s (**Fig. 1G; S2A**). Lastly, the role of cDC1s was assessed in the PyMT breast cancer model (**Fig. S1H**), in which cDC1 deletion has been previously reported to enhance the growth of mammary fat pad (mfp) tumors (15). While this could be reproduced in our study (**Fig. 1I**), strikingly, cDC1 deletion had an opposite effect on the intracranial PyMT tumor growth, both in the intracranial tumor implantation model (**Fig. 1H, I; S2B**) and in hematogenous BrM model where cancer cells are injected into the internal carotid artery (**Fig. 1J, K; S2C-E**).

**Fig. 1.**
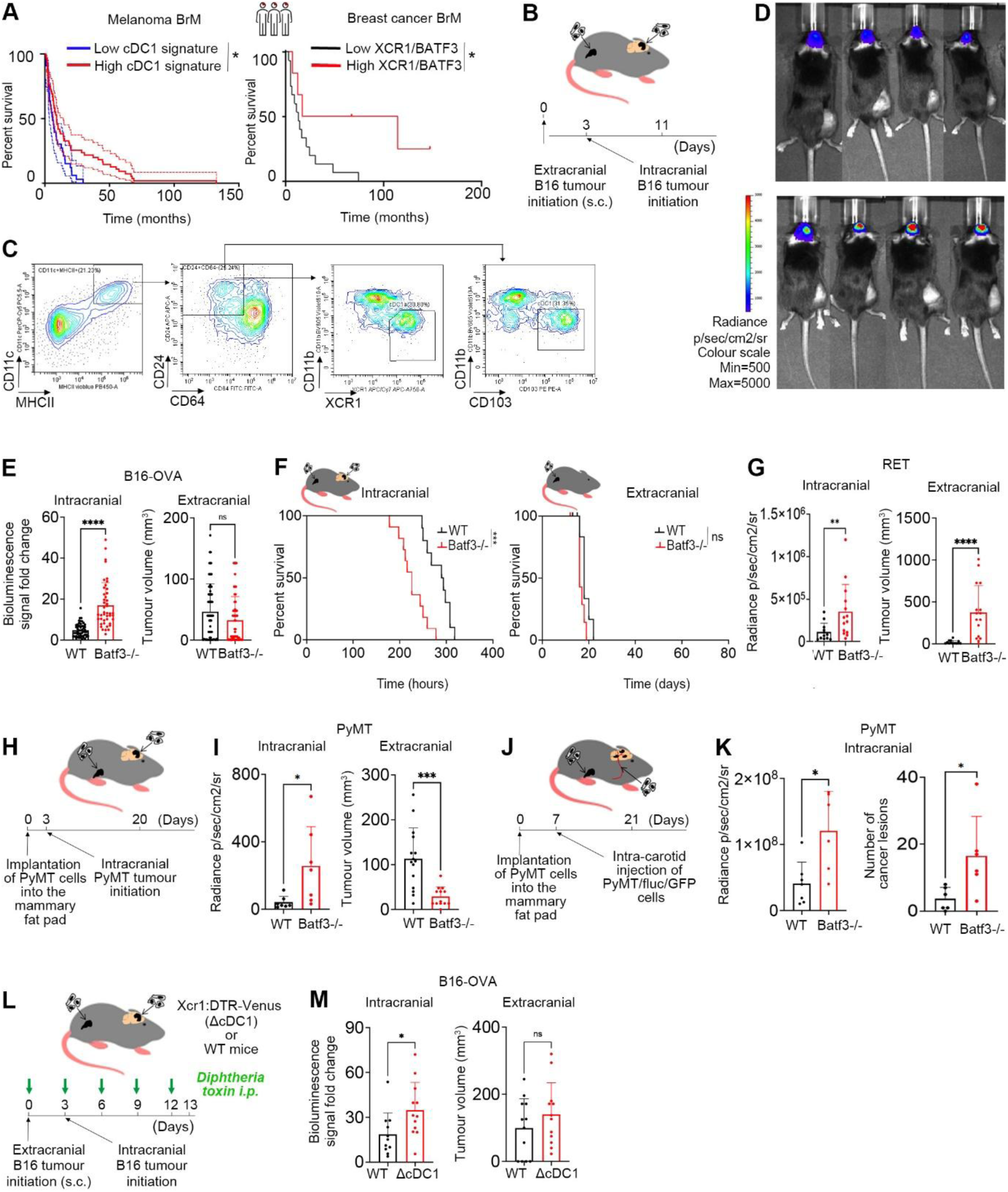
Anti-tumorigenic role of cDC1s in brain metastases. **(A)** Survival from craniotomy in melanoma patients with BrM in group with high (n = 55) and low expression (n = 33) of cDC1 signature **(left);** BrM-specific survival in breast cancer patients with BrM with high (n = 5) and low (n = 16) cDC1 signature **(right)**. **(B)** Two-site B16-OVA model: implantation of B16-OVA cells subcutaneously (extracranially) (4x10^5 cells) and B16-OVA/Fluc cells intracranially (1x10^4 or 1x10^5 cells), with tumor analysis on day 11. **(C)** Gating strategy for identification of cDC1s within CD45+ population by flow cytometry. **(D)** Representative bioluminescence images of the two-site B16-OVA model on day 10. **(E)** Quantification of intracranial tumor burden (fold-change in bioluminescence signal intensity between days 10 and 6; 6 pooled experiments: n = 15/6/11/8/7/7 WT, n = 8/6/7/7/6 Batf3-/-) and extracranial tumor volume (day 11; 9 pooled experiments: n = 6/15/8/7/8/7/7/7/6 WT, n = 7/7/7/6/7/7/7/6/7 Batf3-/-) in B16-OVA two-site model. **(F)** Intracranial tumor-dependent survival in the two-site B16-OVA model (n = 13 WT, n = 8 Batf3-/-) and extracranial tumor dependent survival in the subcutaneous only B16-OVA tumor model (n = 7 WT, n = 7 Batf3-/-). **(G)** Quantification of intracranial tumor burden (bioluminescence signal intensity) and extracranial tumor volume in the RET two-site melanoma model on day 14 (2 pooled experiments: n = 6/8 WT, n = 7/9 Batf3-/-). **(H)** Two-site PyMT breast cancer model (direct intracranial tumor implantation): timeline for implantation of PyMT cells into the mammary fat pad (mfp; extracranially) (2x10^5 cells) and PyMT/Fluc cells intracranially (1x10^5 cells), with tumor harvest on day 20. **(I)** Quantification of Intracranial tumor growth (fold-change in bioluminescence signal intensity between days 20 and 7) and extracranial tumor volume (day 20) in the PyMT two-site model (intracranial: n = 7; extracranial: n = 8). **(J)** Hematogenous two-site PyMT model: timeline for implantation of PyMT cells into the mfp (2x10^5 cells) and PyMT/Fluc/GFP cells into the internal carotid artery (1x10^5 cells), with mfp tumor and whole brain harvest on day 21. **(K)** Hematogenous PyMT two-site model; **Left:** quantification of intracranial tumor burden (bioluminescence signal intensity) on day 21 (n = 7 WT, n = 6 Batf3-/-); **Right:** quantification of intracranial GFP+ cancer lesions within every 4^th^ coronal section of the whole brain (n = 6 WT, n = 6 Batf3-/-). **(L)** Two-site B16-OVA model in Xcr1:DTR-Venus and control mice (WT littermates) indicating injection of Diphtheria toxin (DT) every 3 days, with B16-OVA melanoma cells implanted subcutaneously (4x10^5 cells) and B16-OVA/Fluc cells intracranially (1x10^4 cells), and tumor harvest on day 13. **(M)** Quantification of intracranial tumor burden (bioluminescence signal intensity between days 12 and 7) and extracranial tumor volume (day 12) in the B16-OVA two-site model in WT and Xcr1:DTR-Venus mice (n = 13). Significant differences in **E**, **G**, **I** (intracranial), **K** and **M** (extracranial) were determined by two-tailed Student’s T-test with unequal variance. Significant differences in **E**, **G, I** (extracranial) and **M** (intracranial) were determined by Mann Whitney Test. Significant differences in **A** and **F** were determined by Log rank Test (* < 0.05; ** < 0.01; *** < 0.005; **** < 0.001).

Xcr1:DTR-Venus knock-in mice, in which a fusion protein consisting of Venus fluorescence protein and Diphtheria toxin receptor (DTR) is expressed under the control of the endogenous Xcr1 promoter (16), were used as an additional cDC1 depletion model. Unlike in Batf3-/- mice, where cDC1s lack from birth, in Xcr1:DTR-Venus mice cDC1s were depleted through Diphtheria toxin (DT) injections from the time of extracranial tumor initiation onwards (**Fig. 1L; S2F, G**). In line with the Batf3-/- model, intracranial B16-OVA tumor burden was significantly higher in Xcr1:DTR-Venus as compared to the WT mice, while extracranial tumor burden was not significantly affected following cDC1 depletion (**Fig. 1M**).

In summary, our data demonstrated that while the role of cDC1s at extracranial sites is variable, their role in the brain is consistently anti-tumorigenic independent of the cancer cell model, route of cancer cell injection and cDC1 depletion model, with patient data supporting the association of improved outcomes with higher cDC1 gene signatures in BrM, thus collectively establishing a key role for cDC1s in anti-tumor immune responses in BrM.

### Enhanced activation of intracranial cDC1s is underpinned by type-I-interferon pathway

cDC1s, but not cDC2s, were more abundant within intracranial than extracranial tumors in B16-OVA and PyMT models (**Fig. 2A, B**; **S2H, I**). Both cDC populations were significantly lower in their proportions in the intracranial (cervical) as compared to the extracranial (inguinal) TDLNs (**Fig. 2C**), potentially suggesting a less efficient migration of cDCs from tumors within the brain to the TDLNs.

**Fig. 2.**
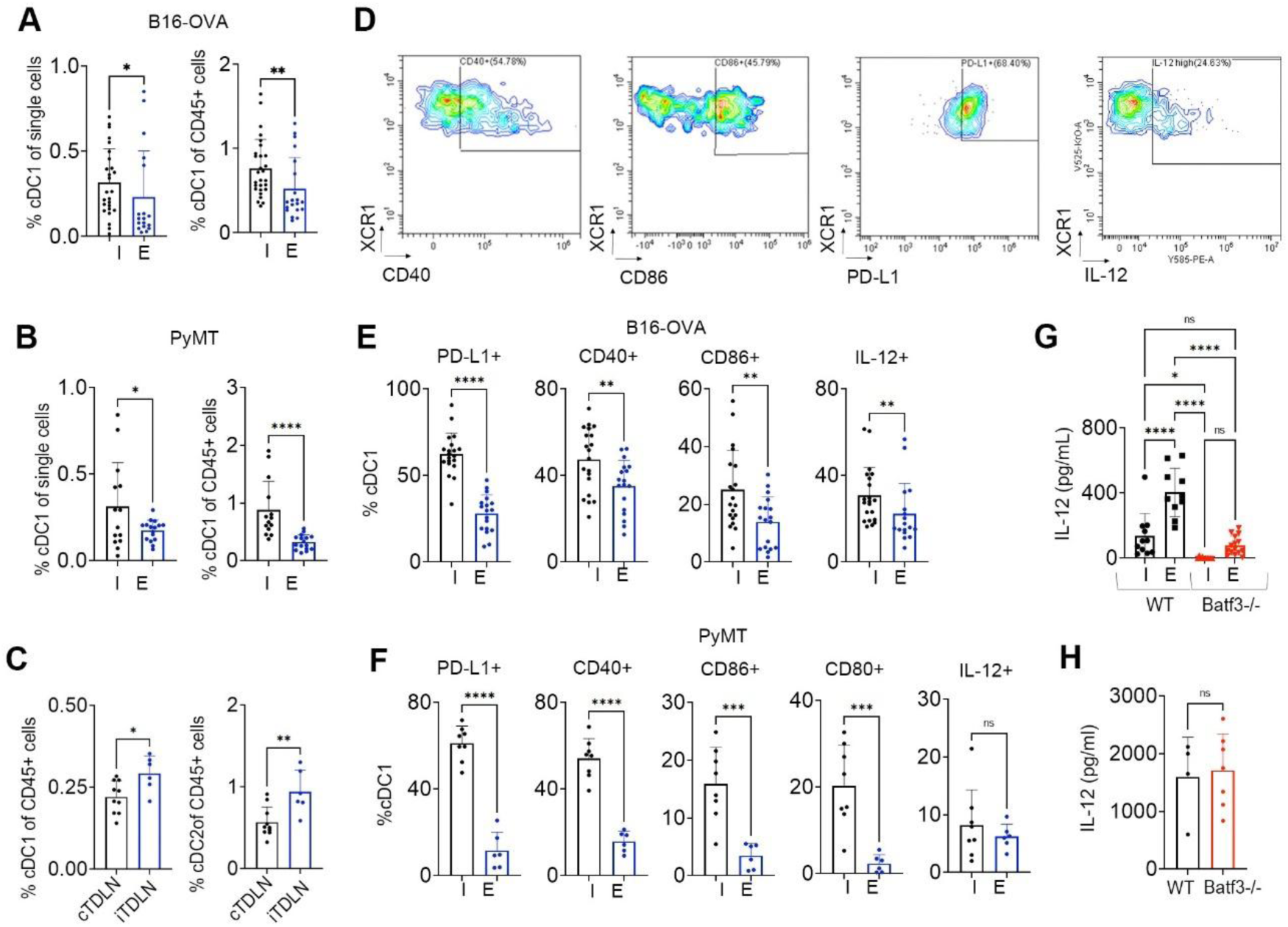
Higher expression of activation markers and IL-12 in intracranial as compared to extracranial cDC1s. (A, **B)** Percentages of cDC1s within single cells (**left**) or within CD45+ cells (**right**) as quantified by flow cytometry within intracranial (I) and extracranial (E) tumors. **(A)** B16-OVA two-site model (4 pooled experiments; Intracranial: n = 7/7/6/6; extracranial: n = 7/4/7/1); **(B)** PyMT two-site model, direct intracranial tumor implantation (2 pooled experiments; Intracranial: n = 8/6; extracranial: n= 7/9). **(C)** Percentages of cDC1s and cDC2s within CD45+ cells quantified by flow cytometry within cervical (c) and inguinal (i) TDLNs in the B16-OVA two-site model (n = 10 cTDLNs; n = 6 iTDLNs). **(D)** Representative flow cytometry plots of cDC1s within intracranial B16-OVA tumors. **(E, F)** Percentages of intra-tumoral PD-L1+, CD40+, CD86+, and IL-12+ cDC1s as quantified by flow cytometry; **(E)** Two-site B16-OVA model (3 pooled experiments; Intracranial: n = 7/7/6; extracranial: n = 8/4/6). **(F)** Two-site PyMT model, direct intracranial tumor implantation (n = 8 intracranial; n = 6 extracranial). **(G)** Quantification of IL-12 protein in tumor supernatants in the two-site B16-OVA model by ELISA. IL-12 concentration (pg/ml) was normalized to tumor weight (2 pooled experiment; Intracranial: n = 3/8 WT, n = 4/8 Batf3-/-; extracranial: n = 6/4 WT, n = 7/8 Batf3-/-). **(H)** Quantification of IL-12 protein in serum in the two-site B16-OVA model by ELISA (n = 4 WT, n = 7 Batf3-/). Significant differences in **B (left), C, F** and **H** were determined by two-tailed Student’s T-test with unequal variance, in **A (left** and **right), B (right),** and **E** by Mann Whitney Test, and in **G** by ordinary one-way ANOVA (* < 0.05; ** < 0.01; *** < 0.005; **** < 0.001).

A significantly higher proportion of intracranial as compared to extracranial cDC1s expressed CD40, CD86, PD-L1 and IL-12 in the B16-OVA (**Fig. 2D, E; Fig. S2J**) and PyMT models (**Fig. 2F**), with only a trend seen for IL-12 in the latter. However, total intra-tumoral IL-12 protein levels were significantly lower in intracranial than extracranial tumors and significantly reduced at both sites in the absence of cDC1s (**Fig. 2G**), despite comparable serum IL-12 levels in WT and Batf3-/- mice (**Fig. 2H**). While a substantial amount of IL-12 remained within extracranial Batf3-/- tumors, IL-12 was abolished in intracranial Batf3-/-tumors (**Fig. 2G**), suggesting that cDC1s are the main and potentially only source of IL-12 in the brain, partially explaining the stronger and more consistent dependence of the intracranial as compared to the extracranial anti-tumor immune responses on cDC1s.

Profiling of cDC1s isolated from B16-OVA tumors by mRNAseq revealed site-specific cDC1 molecular profiles (**Fig. 3A; S3A**). Gene set enrichment analysis (GSEA) using GO database revealed a significant downregulation of pathways related to the positive regulation of ERK1/2 and MAPK, and MyD88-dependent toll-like receptor (TLR) signaling pathway in intracranial cDC1s (**Fig. 3B; S3B**), with *MyD88*, *Tlr1*, *4, 5* and *8* among the downregulated genes (**Fig. 3D, F**). Simultaneously, pathways related to IFNα production and response were significantly upregulated in intracranial cDC1s (**Fig. 3C**; **S3C**), with upregulation of genes encoding *Ifnar1, Ifnar2, Tlr3* and *Irf7* transcription factor known to be upstream of IFNα production (**Fig. 3E, F**). “Antigen processing and presentation”, with a significantly increased expression level of *Erap1,* was among the non-significantly enriched GO terms in intracranial cDC1s (**Fig. S3D**). Expression levels of *Ccr7*, encoding chemokine receptor essential for DC migration to TDLNs, and *Cxcl9/10* encoding T cell-attracting chemokines, were similar between intracranial and extracranial cDC1s (data not shown).

**Fig. 3.**
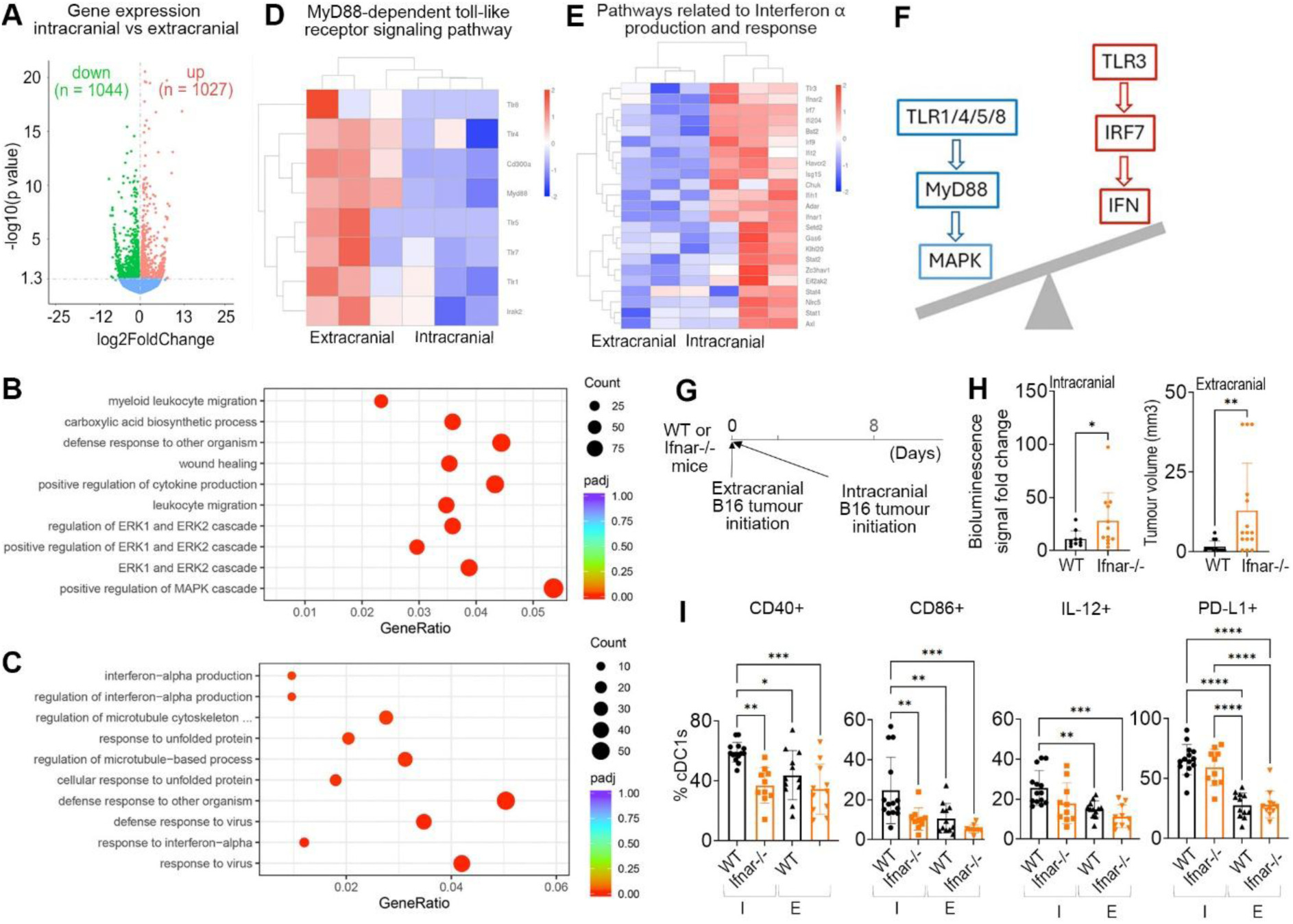
Tumor site-specific cDC1 molecular profiles and type-I-interferon driven upregulation of costimulatory molecules/activation markers in intracranial cDC1s. **(A)** Volcano plot showing differentially expressed genes between cDC1s isolated from intracranial and extracranial B16-OVA tumors, with upregulated genes in red (n = 1027), downregulated genes in green (n = 1044), and genes with unaltered expression in blue (n = 25015). **(B)** Top 10 GO Biological Processes significantly (FDR≤0.05) differentially enriched between intracranial and extracranial cDC1s. **(C)** Top 10 GO Biological Processes significantly upregulated in intracranial versus extracranial cDC1s. **(D, E)** Heat maps showing differentially expressed genes within the “MyD88 dependent TLR signaling pathway” (D) and “Interferon alpha signaling pathway” (E), identified as significantly enriched terms following GSEA (GO database) on genes differentially expressed between intracranial and extracranial cDC1s (FDR≤0.05). **(F)** A scheme summarizing the key genes/pathways downregulated (blue) and upregulated (red) in intracranial cDC1s involved in increased type-I-interferon signaling. **(G)** Two-site B16-OVA model in WT and Ifnar-/- mice: implantation of B16-OVA melanoma cells subcutaneously (extracranially) (4x10^5 cells) and B16-OVA/Fluc cells intracranially (1x10^5 cells), with tumor harvest on day 8. **(H)** Quantification of intracranial tumor burden (fold-change in bioluminescence signal intensity between days 7 and 3) and extracranial tumor volume (day 7) in the B16-OVA two-site model (2 pooled experiments; WT: n = 4/7 intracranial, n = 8/7 extracranial; Ifnar-/-: n = 4/8 intracranial, n = 8/8 extracranial). **(I)** Percentages of CD40+, CD86+, IL-12+ and PD-L1+ cDC1s as quantified by flow cytometry within intracranial (I) and extracranial tumors (E) in the B16-OVA model (2 pooled experiments; WT: n = 7/7 intracranial, n = 8/4 extracranial; Ifnar-/-: n = 6/4 intracranial, n = 4/7 extracranial. Significant differences in **H** were determined by Mann Whitney test and in **I** by Kruskal-Wallis test (for PD-L1) or by ordinary one-way ANOVA (for CD40, CD86, IL-12) (* < 0.05; ** < 0.01; *** < 0.005; **** <0.001).

Because type-I-interferons are known to promote cDC1 ability to upregulate costimulatory molecules and activate CD8+ T cells (17), we hypothesized that differences in this pathway may underpin the tumor site-specific cDC1 activation state. In line with the well-known pleiotropic anti-tumor effects of type-I-IFN (18), we observed a significantly higher intracranial and extracranial tumor burden in Ifnar-/- as compared to the WT mice (**Fig. 3G, H**). However, *Ifnar* deletion resulted in a significantly reduced proportion of cDC1s expressing CD40 and CD86, and a tendency towards reduced proportion of IL-12+ cDC1s within intracranial tumors, without affecting extracranial cDC1s (**Fig. 3I**), suggesting that the enhanced intracranial cDC1 activation is at least partially driven by type-I-IFN.

### Tumor site-specific impact of cDC1 deletion on tumor-infiltrating immune cells

We next analyzed the impact of cDC1 deletion on other immune cells (**Fig. S4A**). In the B16-OVA model, the percentages of CD45+ cells, CD8+ and CD4+ T cells were significantly reduced at both tumor sites in Batf3-/- mice (**Fig. 4A, B; Fig. S4B**), with a stronger reduction observed in intracranial tumors (**Fig. 4C, D; Fig. S4C**), while T cells within TDLNs remained unaffected (**Fig. S4D, E**). cDC1 deletion also significantly reduced CD8+ T cell proportions in intracranial tumors in the RET melanoma model, PyMT direct intracranial tumor implantation model and PyMT hematogenous model, with only a non-significant tendency towards reduced CD8+ T cells in extracranial RET tumors and no effect in extracranial PyMT tumors (**Fig. 4E-G**). Thus, cDC1 deletion more prominently reduced the abundance of CD8+ T cells within intracranial than extracranial tumors in a model independent manner, reiterating a stronger reliance of intracranial anti-tumor immunity on cDC1s.

**Fig. 4.**
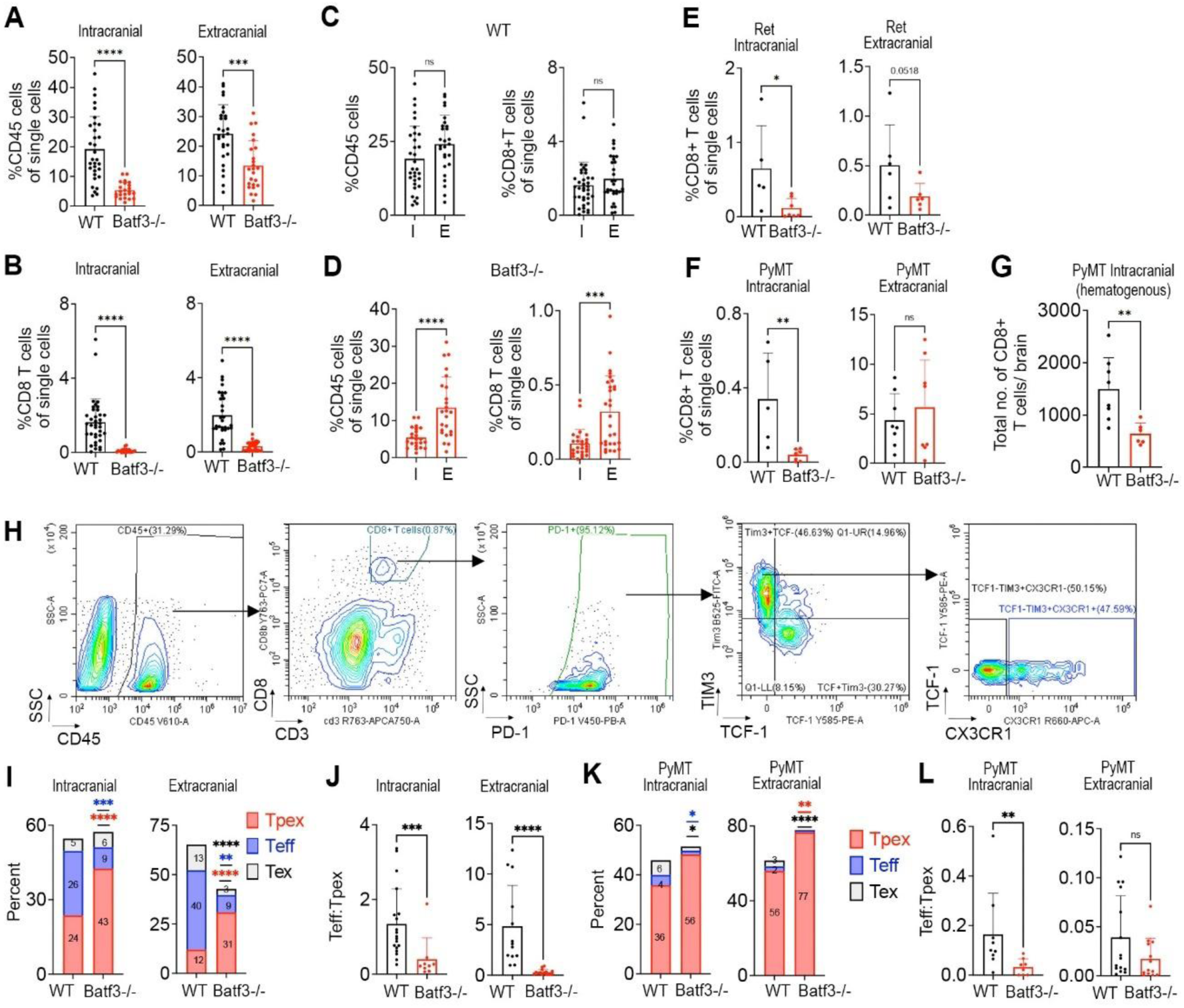
cDC1s are critical for the maintenance of the intra-tumoral CD8+ T cell pool and efficient conversion of progenitor exhausted T cells to effectors. **(A, B**) Percentages of intra-tumoral CD45+ cells (**A**) and CD8+ T cells (**B**) within single cells in the two-site B16-OVA model, as quantified by flow cytometry (WT (5 pooled experiments): n = 5/7/7/6/7 intracranial, n = 5/3/10/6/3 extracranial; Batf3-/- (4 pooled experiments): n = 3/7/7/7 intracranial, n = 4/4/9/7 extracranial. **(C, D)** Data from A and B were replotted for direct comparison of percentages of CD45+ cells and CD8+ T cells between the intracranial (I) and extracranial (E) tumors in the WT (C) and Batf3-/- mice (D). **(E, F)** Percentages of intra-tumoral CD8+ T cells within single cells in the two-site Ret melanoma model (intracranial: n = 5 WT, n = 7 Batf3-/-; extracranial: n = 6 WT, n = 6 Batf3-/-) (E) and PyMT model with direct intracranial tumor implantation (n = 5 WT, n = 8 Batf3-/-) (F). **(G)** Total number of CD8+ T cells per brain in the hematogenous two-site PyMT model as quantified by flow cytometry (n = 8 WT, n = 6 Batf3-/-). **(H)** Representative flow cytometry plots of intracranial B16-OVA tumors showing the gating strategy for identification of the exhausted T cell subsets Tpex (TCF-1+TIM3-), Teff (TCF-1-CX3CR1+) and Tex (TCF-1-CX3CR1-) within CD45+CD3+CD8+PD1+ T cells. **(I, K)** Percentages of intra-tumoral Tpex, Teff, and Tex within CD8+PD1+ T cells in the two-site B16-OVA model (I) and PyMT model with direct intracranial tumor implantation) (K). **(J, L)** Ratio of Teff to Tpex in the two-site B16-OVA model (J) and PyMT model with direct intracranial tumor implantation (O). Data in **I** and **J** (B16-OVA model) were pooled from 3 independent experiments; intracranial: n = 7/7/5 WT, n = 1/4/4 Batf3-/-; extracranial: n = 6/5/3 WT, n = 6/5/7 Batf3-/-. Data in **K** and **L** (PyMT model) were pooled from 2 independent experiments; intracranial: n = 5/5 WT, n = 4/4 Batf3-/-; extracranial: n = 8/8 WT, n = 8/7 Batf3-/-. Significant differences in **D (left)**, **E, F, G, I (left, Tpex and Teff; right, Teff),** and **K** were determined by two-tailed Student’s T-test with unequal variance. Significant differences in **A-C, D (right), I (right Tpex, Tex), J** and **L** were determined by Mann Whitney Test (* < 0.05; ** < 0.01; *** < 0.005; **** < 0.001).

### cDC1 population is required for the differentiation of exhausted CD8+ T cell subsets

Conversion of Tpex to Teff with cytotoxic functions within tumors is critical for anti-tumor immune responses. As the dynamics of exhausted CD8+ T cells in BrM and their dependence on cDC1s are unknown, we investigated this in our study (**Fig. 4H; S5A**). Intracranial B16-OVA tumors displayed significantly lower Teff numbers (**Fig. S5B**), lower Teff and Tex proportions within CD8+ T cells (**Fig. S5C**), and thus lower Teff/Tpex ratio as compared to extracranial tumors (**Fig. S5D**). The intracranial Tex numbers were very low, and except for this population, all exhausted T cell subsets were significantly reduced in Batf3-/- as compared to the WT mice at both tumor sites (**Fig. S5E**). The extent of reduction, however, differed between the subsets and most drastically affected Teff population, with 58-fold and 36-fold reduction in cell numbers in intracranial and extracranial tumors, respectively (**Fig. S5E**), leading to increased Tpex and reduced Teff proportions within CD8+ T cells in Batf3-/- mice (**Fig. 4I**), and thus significantly reduced Teff/Tpex ratio in the absence of cDC1s at both tumor sites (**Fig. 4J**).

In the PyMT breast cancer model, CD8+ T cells and the three subsets displayed a lower abundance in intracranial than extracranial tumors (**Fig. S5F, G**). Tpex proportion within CD8+PD1+ T cells was lower, while Teff proportion, as well as Teff/Tpex ratio were significantly higher in intracranial versus extracranial PyMT tumors (**Fig. S5H, I**), contrasting the observations in melanoma model. While cDC1 deletion had no effect on Tpex numbers, Teff numbers were significantly reduced in Batf3-/- versus WT mice at both tumor sites (**Fig. S5J**). Importantly, similarly to the melanoma model, cDC1 deletion reduced the proportion of Teff within intracranial CD8+PD1+ T cells and reduced proportion of Tex at both tumor sites, with a concomitant increase in Tpex proportions, which reached statistical significance in extracranial tumors only (**Fig. 4K**). These changes resulted in a significantly reduced Teff/Tpex ratio in the intracranial PyMT tumors and a similar trend in extracranial tumors (**Fig. 4L**). Thus, while contrasting tumor site-specific differences in the proportions of CD8+ T cells and their subsets were observed in melanoma and breast cancer models, the impact of cDC1 deletion was model-independent, with a drop in Teff/Tpex ratio at both tumor sites, demonstrating that tumors in the brain also rely on cDC1s for Tpex-to-Teff conversion.

### cDC1 deletion causes a stronger impairment of intracranial than extracranial ICB efficacy

We next analyzed the role of cDC1s in the efficacy of a combined anti-PD-1 / anti-CTLA-4 (PC) therapy, chosen due to its superior clinical efficacy over monotherapies in melanoma BrM (2, 3). While PC therapy significantly extended both intracranial and extracranial tumor-dependent survival in WT and Batf3-/- mice, this extension was significantly smaller in the Batf3-/- mice (**Fig. 5A-D**). PC blockade also failed to reduce intracranial and extracranial tumor burden in Batf3-/- mice, in contrast to a significant reduction in WT mice (**Fig. 5E-G**). Moreover, while PC blockade increased the percentage of CD45+ cells and CD8+ T cells in intracranial and extracranial tumors in the WT and Batf3-/- mice (**Fig. 5H**), the proportion of both populations was significantly lower in treated Batf3-/- as compared to the treated WT mice (**Fig. 5H**), demonstrating that cDC1s are important for the efficacy of PC therapy at both tumor sites. However, while in the absence of cDC1s the PC blockade extended the extracranial tumor-dependent survival by 73.6% (16.3±2.05 days for IgG and 28.3±19.7 days for PC group) (**Fig. 5D**), the intracranial tumor-dependent survival was extended by only 13.3% (227±29.0 hours for IgG and 257±47.0 hours for PC group) (**Fig. 5B**). Moreover, a direct comparison between the two tumor sites in the treated Batf3-/- mice revealed a significantly lower proportion of CD45+ cells and a trend towards lower percentage of CD8+ T cells in intracranial tumors (**Fig. 5J**), in summary demonstrating a stronger reliance of intracranial than extracranial anti-tumor immunity on cDC1s for the efficacy of PC therapy.

**Fig. 5.**
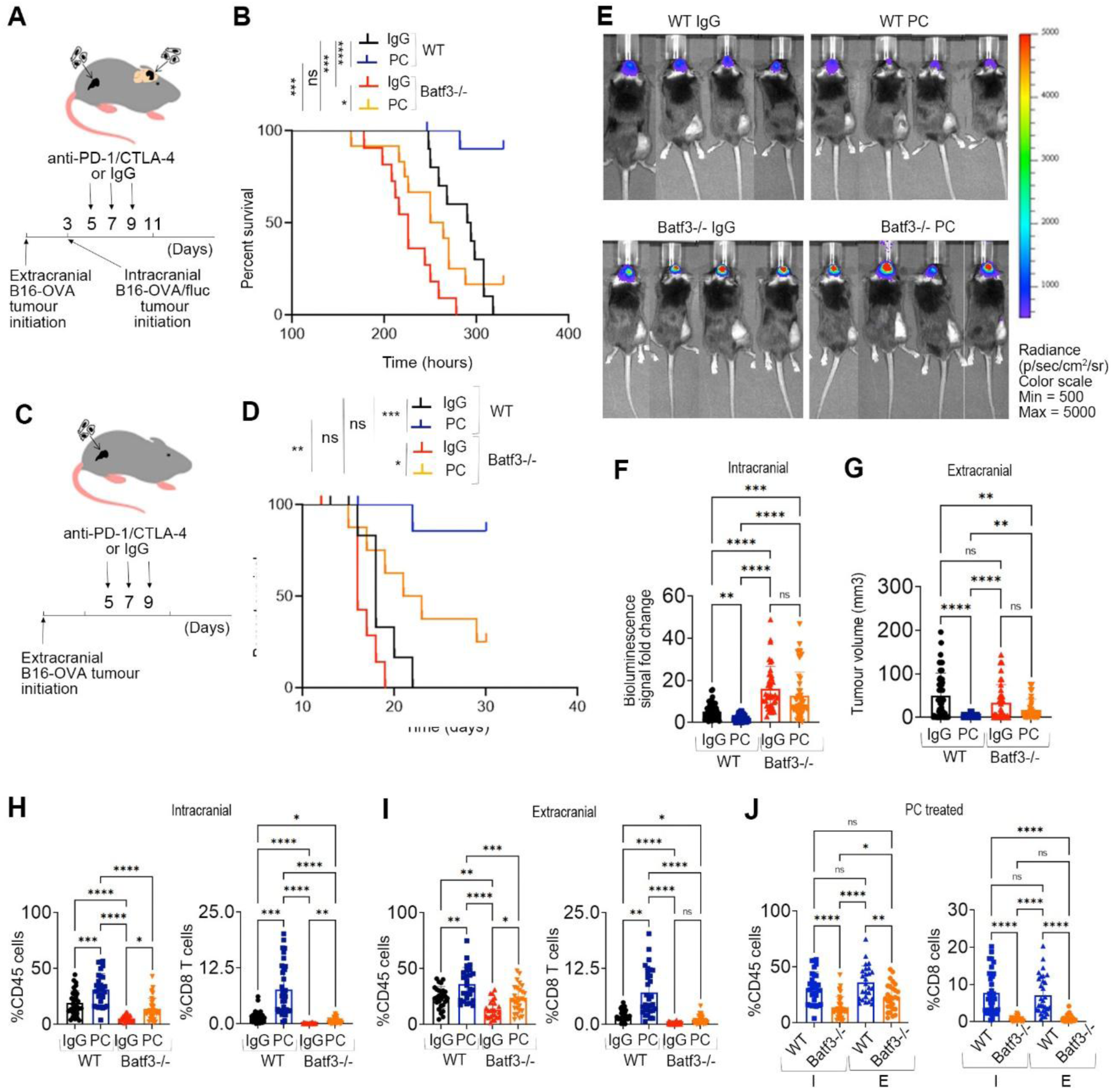
cDC1 deletion more severely impairs the intracranial than extracranial efficacy of the combined PD-1/CTLA-4 blockade. **(A, C)** Experimental timelines for the two-site B16-OVA model (A) and the subcutaneous only tumor model (C), with implantation of B16-OVA cells subcutaneously (extracranially) (4x10^5 cells) and B16-OVA/Fluc cells intracranially (1x10^4 cells), and i.p. administration of therapeutic antibodies (anti-CTLA-4 plus anti-PD-1 (PC), or IgG control). **(B, D)** Intracranial tumor-dependent survival in the two-site B16-OVA model (n = 11 WT IgG, n = 11 WT PC, n = 11 Batf3-/- IgG, n = 12 Batf3-/- PC) (B) and the extracranial tumor dependent survival in mice bearing only subcutaneous tumors (n = 7 WT IgG, n = 8 WT PC, n = 7 Batf3-/- IgG, n = 8 Batf3-/- PC) (D). **(E)** Representative bioluminescence images of mice in the two-site B16-OVA model on day 10. **(F, G)** Quantification of intracranial tumor growth (fold-change in bioluminescence signal intensity between days 10 and 6) (F) and extracranial tumor volume (day 11) (G) in the two-site B16-OVA model. Intracranial: data pooled from 6 experiments for WT IgG, WT PC, and Batf3-/-PC, and from 5 experiments for Batf3-/- IgG; n = 6/15/8/7/8/11 WT IgG, n = 6/16/8/7/8/11 WT PC, n = 5/7/7/7/11 Batf3-/- IgG, n = 6/9/8/6/8/12 Batf3-/- PC. Extracranial: data pooled from 10 experiments for WT IgG, WT PC, and Batf3-/- PC, and from 9 experiments for Batf3-/-IgG; n = 6/15/8/7/8/7/8/7/7/6 WT IgG, n = 8/16/8/7/8/7/8/7/7/8 WT PC, n = 7/7/7/6/7/7/7/6/7 Batf3-/- IgG, n = 7/9/8/6/8/7/8/7/8/7 Batf3-/- PC. **(H, I)** Percentages of intra-tumoral CD45+ cells and CD8+ T cells within single cells in the two-site B16-OVA model, as quantified by flow cytometry. Data pooled from 5 experiments for WT IgG, WT PC, and Batf3-/- PC, and 4 experiments for Batf3-/- IgG. Intracranial: n = 5/7/7/6/7 WT IgG, n = 4/6/6/6/7 WT PC, n = 3/7/7/7 Batf3-/- IgG, n = 5/7/7/10/7 Batf3-/- PC; extracranial: n = 5/3/10/6/3 WT IgG, n = 3/4/9/6/4 WT PC, n = 4/4/10/7 Batf3-/- IgG, n = 5/4/10/5/4 Batf3-/- PC. **(J)** Data from the PC blockade treated groups from H-I were replotted to allow for a direct comparison of the percentages of CD45+ cells and CD8+ T cells between WT and Batf3-/- mice. Significant differences in **B** and **D** were determined by Log rank Test. Significant differences in **F, G, H (right), I (right),** and **J (right)** were determined by Kruskal-Wallis test, and in **H (left), I (left),** and **J (left)** by ordinary one-way ANOVA (* < 0.05; ** < 0.01; *** < 0.005; **** < 0.001).

### cDC1 population is required for the PC blockade-driven expansion of CD8+ Teff in brain metastases

We next asked which exhausted CD8+ T cell subsets are expanded the most by PC therapy. Only Teff population was significantly increased in numbers in intracranial B16-OVA tumors following PC blockade, with a concomitant trend towards increase in Tex numbers (**Fig. 6A**). This resulted in a significantly increased Teff proportion within CD8+ T cells and a trend towards reduced Tpex proportion following therapy (**Fig. 6B**), suggesting that PC blockade enhances Tpex-to-Teff conversion in BrM.

**Fig. 6.**
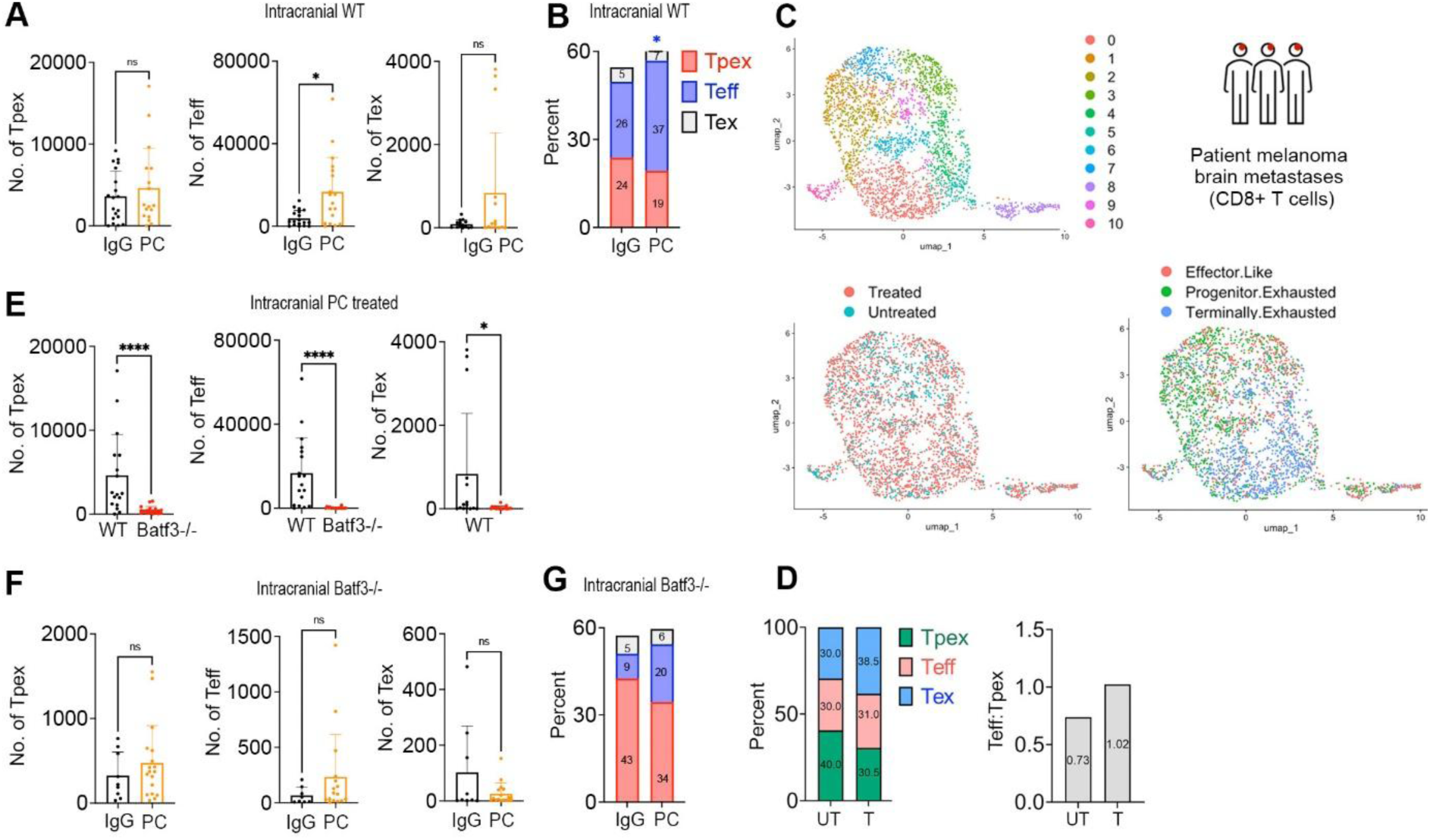
PC blockade driven Teff expansion requires cDC1s. **(A, B)** Numbers of Tpex, Teff and Tex per 100,000 single cells (A) and their proportions within CD8+ T cells (B) in intracranial tumors in the two-site B16-OVA model in WT mice receiving PC blockade or IgG control, as quantified by flow cytometry (3 pooled experiments; n = 5/7/7 IgG, n = 4/7/7 PC). (B) Exhausted T cell subsets within CD8+ T cells from the human melanoma brain metastases scRNAseq dataset by Alvarez-Brackenridge et al. (19) were identified using Tpex, Teff and Tex signatures extracted from Miller et al. (10). Data set included cells from 7 immunotherapy naïve and 23 immunotherapy treated samples (n = 4 PD-1 blockade, n = 17 combined PD-1/CTLA-4 blockade) obtained from 26 patients. UMAPs displaying clustering of CD8+ T cells (**top left**), annotation of CD8+ T cells from treated and untreated samples (**bottom left**), and annotation of exhausted T cell subsets (**bottom right**) are shown. **(D)** Data from C: quantification of the proportions of exhausted CD8+ T cell subsets (**left**) and Teff/Tpex ratio (**right**) in human melanoma brain metastases from immunotherapy naïve (untreated; UT) and immunotherapy treated patients (T). **(E)** Data from A and F were replotted for direct comparison of T cell numbers between PC-treated WT and PC-treated Batf3-/- mice. **(F-G)** Numbers of Tpex, Teff and Tex per 100,000 single cells (F) and their proportions within CD8+ T cells (G) in intracranial tumors in the two-site B16-OVA model in the Batf3-/- mice receiving PC blockade or IgG control, as quantified by flow cytometry (3 pooled experiments; n = 4/4/1 Batf3-/- IgG, n = 6/8/8 Batf3-/- PC). Significant differences in **A, B, E** and **F** were determined by Mann Whitney Test (* < 0.05; ** < 0.01; *** < 0.005; **** < 0.001).

To investigate whether this is also the case in patients, the signatures of exhausted T cell subsets extracted from Miller et al. (10) were applied to CD8+ T cells extracted from the previously published scRNAseq dataset of human melanoma BrM, composed of samples collected following ICB and immunotherapy-naïve samples (19) (**Fig. 6C**). ICB reduced the proportion of Tpex and increased the proportion of Tex, thereby increasing the Teff/Tpex ratio (**Fig. 6D**). This suggested that, in line with our preclinical data, PC blockade in patients promotes differentiation of Tpex in BrM towards Teff/Tex.

We next focused on the role of cDC1s in the PC blockade-induced changes in the dynamics of exhausted CD8+ T cell subsets. Similarly to the untreated mice, in the presence of PC blockade all three exhausted T cell subsets were significantly reduced in numbers in Batf3-/- as compared to the WT mice (**Fig. 6E**). Notably, the increase in Teff numbers and Teff proportions that were observed following PC blockade in the WT mice were no longer significant in the absence of cDC1s (**Fig. 6F, G**), demonstrating that PC blockade-induced Teff augmentation is at least partially cDC1-dependent. This demonstrated that the Tpex-to-Teff conversion and/or Teff expansion in BrM relies on cDC1s not only during tumor establishment, but also for a successful therapeutic response to ICB.

## Discussion

While cDC1s and their role in immunotherapy have been studied in the context of extracranial tumors and primary brain tumors (5–7, 20), their role in BrM remained understudied. Using two-site BrM models, which mimic the presence of clinically observed extracranial cancer lesions and clinical intracranial responses to ICB (13), we here demonstrate that the cDC1 population is consistently critical for the control of tumor growth in the brain in melanoma and breast cancer models, despite its variable and model-dependent role in extracranial tumors, and its reported lack of involvement in tumor growth in malignant primary brain tumor models (20). These findings are supported by a positive correlation between cDC1 gene signature and BrM-specific survival in melanoma and breast cancer patients.

Our study demonstrates a higher and more consistent dependence of anti-tumor immune responses on cDC1s in intracranial as compared to extracranial tumors, during tumor establishment and during ICB therapy, and identifies potential mechanisms underpinning these observations. In comparison to extracranial cDC1s, intracranial cDC1s demonstrated higher abundance and significantly higher expression of costimulatory molecules, they appeared to be the sole producer of IL-12 within intracranial tumors, and their deletion more severely impaired the abundance of intra-tumoral CD8+ T cells. Overall, this suggests that T cells in the brain heavily rely on cDC1s for co-stimulation, activation and/or recruitment, while in extracranial tumors these processes are likely to be supported by additional antigen presenting cells, as demonstrated for cDC2s in a breast cancer model (15). This sole reliance on cDC1s could partially contribute to a higher threshold for induction of anti-tumor immune responses in the brain, adding to the immune specialized characteristics of the brain microenvironment. At the molecular level, the higher cDC1 activation in the brain as compared to the extracranial tumors seems to be at least partially driven by stronger activation of the type-I-interferon pathway, and could be underpinned by higher expression of the type-I-interferon receptor (IFNAR) on intracranial as compared to extracranial cDC1s, site-specific differences in local type-I-interferon concentrations, and/or site-specific cDC1 TLR repertoire, leading to the downregulation of the MyD88/MAPK pathway and upregulation of IRF7/IFN pathway in intracranial cDC1s. These site-specific differences in molecular profiles could be due to the different origin of intracranial and extracranial cDC1s or imposed by different tumor microenvironments.

While cDC1 deletion didn’t affect the abundance of total CD8+ T cells in TDLNs, we did not investigate whether it altered Tpex proportions. cDC1s have been shown to be required for Tpex maintenance within the extracranial TDLNs (9), but it remains to be determined in future studies whether this differs in the intracranial TDLNs, considering the reported tolerogenic nature of the immune responses in the LNs at this location (21). Focusing on tumors, our study demonstrated that, in the absence of therapy as well as in the context of PC blockade, cDC1s are involved in the Tpex-to-Teff conversion and/or Teff expansion within intracranial tumors. Importantly, PC blockade-induced shift towards more differentiated CD8+ T cell subsets (Teff and Tex) was also observed in melanoma patient BrM, supporting our preclinical findings.

Notably, the Teff/Tpex ratio in tumors has been shown to increase over time (10). Therefore, the tumor site-specific differences in Teff/Tpex ratio in the B16-OVA model could have been caused by a shorter time between tumor implantation and the endpoint in the brain as compared to the extracranial tumors. Despite these constraints in interpreting our findings, different dynamics of CD8+ T cell subsets at the two sites cannot be excluded. Interestingly, a study by Sudmeier et al. (22) suggested that a large proportion of Tpex population in BrM are non-tumor specific bystander cells, and the T cell receptor overlap between the progenitor and exhausted T cell clusters was low, suggesting that a substantial proportion of tumor antigen-specific Tex are derived from progenitors outside BrM. In line with this, our previous study demonstrated that the presence of extracranial tumor in mice augments the PC blockade-driven infiltration of effector CD8+ T cells into intracranial melanoma tumors, implying that the BrM-infiltrating CD8+ T cells are at least in part activated/expanded outside the brain by extracranial cancer lesions (13). Thus, extracranially activated Teff may contribute to the intracranial Teff pool following PC blockade. While these hypotheses will need to be experimentally validated, this is beyond the scope of the current study focusing on cDC1s.

A recent study demonstrated a key role for CCR7+ conventional DCs in controlling cancer cell growth in leptomeningeal metastases (23). While these leptomeningeal DCs consisted mainly of cDC2 population, it cannot be excluded that cDC1s also play an important role not only in the parenchymal BrM but also in the leptomeningeal BrM, a biologically very distinct tumor entity.

In summary, our study establishes a key role for cDC1s in the control of tumor growth in BrM. Our observations on the high reliance of the intracranial anti-tumor immunity on cDC1s imply that DC therapies focusing on cDC1s, potentially in combination with other immunotherapies, especially those acting to activate type-I-IFN pathway, have a high potential to improve tumor control in the brain.

## Methods

### Animals

6–12-week-old female and male C57BL/6J, *Batf3-/-* (B6.129S(C) *Batf3tm1Kmm*/J; Jax stock 013755), and *Ifnar-/-* mice (B6.129S2-*Ifnar1tm1Agt*/Mmjax Jax stock nm. 032045) were purchased from Charles River Laboratories. Xcr1:DTR-venus mice (16) were provided by the RIKEN BRC through the National BioResource Project of the MEXT, Japan. Mice were housed in special pathogen-free facility in individually ventilated cages with Tecniplast IVC Caging Type GM500, with 3R’s bedding material and an automatic 12-hour light / 12-hour dark cycle with temperature maintained at 19-23°C. Food type was SDS (Special Diet Services) CRM diet. Sizzle nest and plastic domes were used as environmental enrichment. Mice were acclimatized for at least 7 days prior to any procedures. Only female mice were used in the breast cancer model, as this cancer type occurs predominantly in females, while both sexes were used in melanoma models. Similar findings were observed for both sexes.

### In vivo models

All in vivo procedures were approved by the University of Leeds Animal Welfare and Ethical Review Committee and performed under the approved United Kingdom Home Office license in line with the Animal (Scientific Procedures) Act 1986 and in accordance with the UK National Cancer Research Institute Guidelines for the welfare of animals (24). In the two-site model, extracranial tumors were implanted into the mammary fat pad (mfp) (breast cancer models) or subcutaneously (melanoma models), followed by intracranial implantation of cancer cells as previously described (13, 25). Briefly, B16-OVA cells (2-6x10^5 cells/20µL of 1:1 Minimum Essential Medium Eagle (MEME) and geltrex) or RET cells (1x10^5 cells/30µL of MEME) were injected subcutaneously on the flank and PyMT cells (2x10^5 cells/10µL of 1:1 PBS and geltrex) were injected into the 4th mammary fat pad to generate extracranial tumors. Three days post-extracranial tumor initiation, 1x10^4 or 1x10^5 B16-OVA/Fluc cells, 1x10^3 RET/fluc cells, or 1x10^5 PyMT/fluc cells in 1µL MEME were stereotactically injected into the striatum (1mm right from the midline, 2mm anterior from bregma, 2.5mm deep) to generate intracranial tumors. In the hematogenous two-site model, PyMT/GFP/Fluc cells (1x10^5 cells in 50µl MEME) were injected into the internal carotid artery 7 days after the injection of PyMT cells into the mfp. Intra-carotid injections were performed as previously described (Rippaus et al., 2016)(Rippaus et al., 2016). Some experiments utilized the one-site subcutaneous model, in which cancer cells were implanted only subcutaneously. In the Xcr1:DTR-Venus mice (homozygous) and the littermate WT mice used as controls, Diphtheria toxin was injected i.p. at 1 µg / mouse for the first injection and at 25 ng / g body weight for all subsequent injections. Mice were monitored daily throughout the experiments and between two and three times per day when approaching the endpoint for neurological symptoms, as well as under-grooming and reduced activity as general signs of compromised health. To exclude the time of day of implantation as a potential confounder, the order of implantation was alternated between the experimental groups. Analgesics were administered prior to surgery and at 24 hours post-surgery.

## Author Contributions

ML conceived the project. ML and FJ designed the experiments. ML, AM, DW and WB supervised the research. FJ, CF, JW, NI, MLaw, ML, GM, AR and DG performed the experiments. FJ, AR and ML analyzed the data. ZH, EJRV and AS performed reanalysis of publicly available gene expression datasets from patient brain metastases. TK generated Xcr1:DTR-Venus mice. FJ wrote the methods and figure legends. ML wrote the abstract, introduction, results, and discussion, and edited the methods and figure legends.

## Acknowledgements

We thank the staff of St. James’s Biological Services Facility, University of Leeds, in particular J. Bilton and D. Evans, and the staff of Biological Services Facility at the University of York, in particular J. Bland and T. Fewlass, for technical support with in vivo experiments. We thank L. Straszynski (University of Leeds), Sukhveer Mann and Karen Hogg (The Bioscience Technology Facility, Department of Biology, University of York) for technical support with flow cytometry. Part of the computational data analyses presented in this work was performed under ARC4, a High-Performance Computing cluster managed by the Advanced Research Computing facility at the University of Leeds, who we’d like to thank. We would like to acknowledge Dr. Atsushi Miyawaki with relation to the use of Venus (27) in the transgenic Xcr1-DTR-Venus mice. We thank Dr. Michael Davies (University of Texas, M.D. Anderson Cancer Center) and the European Genome-Phenome Archive for providing the mRNAseq data of patient brain metastases (11). We acknowledge the use of the scRNAseq data of patient brain metastases (phs002416.v2.p1) obtained from DbGaP, published in Prakadan et al. (28) and Alvarez-Brackenridge et al. (19).

