## Supplemental files for "Conventional dendritic cells type I with an enhanced type-I-IFN signaling underpin anti-tumor immune responses in brain metastases"

### Supporting methods

#### Cell lines and cell culture

B16-OVA cells (29) were kindly provided by Richard Vile, Mayo Clinic. and were cultured in Dulbecco's Modified Eagle Medium (DMEM; SigmaAldrich) supplemented with 10% Fetal Bovine Serum (FBS; Thermofisher), 1% L-glutamine (Sigma) and 1% penicillin / streptomycin (Penstrep; Sigma). PyMT 3503 cells (30) were derived from spontaneous mammary fat pad tumors in MMTV:PyMT mice and were kindly provided by Wolfram Ruf from The Scripps Research Institute, La Jolla, CA. PyMT 3503 cells were cultured in L-15 (Leibovitz) without glucose (Sigma) supplemented with 10% FBS, 1% L-glutamine, 1% Penstrep and insulin (Sigma) at 10ug/ml. RET melanoma cells (31) were kindly provided by Neta Erez, Tel Aviv University, Tel Aviv-Yafo, Israel and were cultured in Roswell Park Memorial Institute Medium (RPMI) 1640 without L-glutamine (Sigma) supplemented with 10% FBS, 1% L-glutamine, 1% Penstrep and 1% Sodium Pyruvate (Gibco). All cells were incubated at 37°C, 5% CO2. For some experiments, cancer cells were used that had been previously stably transduced with firefly luciferase-expressing lentiviral vector pFUW-Fluc (32) or GFP-expressing lentiviral vector pFUGW (33).

#### Quantification of tumor growth

Subcutaneous and mfp tumor growth was quantified by caliper measurement and intracranial tumor growth was quantified by non-invasive bioluminescence imaging using IVIS Spectrum (PerkinElmer). Briefly, mice were injected with 80µL luciferin (Tocris,15mg/ml in PBS) s.c. and imaged after 12 minutes. Living Image software (PerkinElmer) was used for image analysis and quantification of the bioluminescence signal intensity by region of interest (ROI).

#### PC blockade therapy

For experiments with PC blockade, Anti–PD-1 (Bio-X-Cell; clone: RMP1-14; BE0146), anti–CTLA-4 (Bio-X-Cell; clone: 9D9; BE0164), and IgG control antibodies (Bio-X-Cell; Clone: MPC-11; BE0086) were administered intraperitoneally at 200µg in 100µL PBS per mouse. Before treatment, mice were randomized into therapy groups based on the intracranial bioluminescence signals ensuring equal distribution of tumor burden across groups. To minimize the time of day as a variable between experiments all therapeutic treatments took place between 09:00 and 11:00 and within a 30-minute window.

**Immunofluorescence of murine tissue**

After terminal perfusion of mice with PBS, brains were isolated and fixed in 4% paraformaldehyde at 4°C overnight, then transferred into 25% sucrose / 0.1M sodium phosphate buffer and incubated in the fridge until dehydrated. Sectioning of the tissue on the cryotome was performed (40µm sections) and the floating sections were stored in Walter's antifreeze (30% (v/v) ethylenglycol, 30% (v/v) glycerol, and 0.5 M phosphate buffer) at -20°C until use.

Prior to staining, the sections were washed with PBS three times before being blocked in blocking buffer (0.3% Trixton-X-100 (Sigma-Aldrich), 10% goat serum (Thermo Fischer Scientific) or FBS (Thermo fisher) and 10% FcR blocking reagent, mouse (Miltenyi) in PBS) for one hour at room temperature (RT). Sections were then stained with anti-CD103 (R&D Systems, AF1990; 1:100) and anti-MHCII (BD 556999; 1:100) or anti-GFP (Abcam, ab13970; 1:600) diluted in the blocking buffer over night at RT. The samples were washed again in PBS three times and then incubated for 1 hour at RT with the secondary antibodies anti-goat IgG (H+L) Alexa Fluor™ Plus 647 (Invitrogen; A32849 1:500) and anti-rat-Cy3 (Jackson; 712-165-153 1:200) diluted in PBS with 3% bovine serum albumin (BSA), or anti-chicken-FITC (Jackson, 703-096-15; 1:200) diluted in the blocking buffer. DAPI (AdipoGen Life Sciences; 2 µg/mL in PBS) was added for 10 minutes at RT before the final washing in PBS. ProLong Gold Antifade mounting medium (Invitrogen) was then added to each tissue slide and sections covered with a glass coverslip. Sections were imaged using the Widefield

Fluorescent Inverted Microscope Nikon Eclipse Ti-E. QuPath v0.5.1 was used for image analysis and production. Quantification of GFP+ tumor areas was performed blinded.

**Dissociation of Murine Tumors and collection of tumor supernatants for ELISA**

Mice were terminally perfused with PBS and s.c. tumors, mfp tumors, whole brain and LNs were isolated. Brain tumors from the intracranial tumor implantation models were macro dissected from the surrounding brain tissue, mechanically disrupted with scalpels, resuspended in 200 to 1000 µL collagenase solution (3 mg/mL collagenase A (Roche; 10103578001), 250 U/mL hyaluronidase (Sigma; H-3506), 30 U/mL DNAseI (Sigma; 04716728001) in MEME (Sigma; M2279)), and incubated for 15 minutes at 37°C, followed by dissociation through pipetting. The cells were collected by centrifugation at 400 g for 5 minutes at 4°C. The supernatants were transferred into fresh tubes and stored frozen at -80°C for later analysis by ELISA. Cell pellets were resuspended in the incubation buffer (2mM EDTA, 0.5% bovine serum albumin (BSA; Sigma-Aldrich) in PBS) and strained through a 100 µm cell strainer. For analysis of immune cells in the hematogenous two-site model, the whole brain was dissociated as described above, and myelin was removed with the myelin removal beads (Miltenyi) following the manufacturer's protocol. LNs were placed directly into collagenase solution, incubated for 15 minutes at 37°C, mechanically disrupted by pipetting, washed with incubation buffer and strained through a 100 µm cell strainer to obtain a single-cell suspension.

**Flow Cytometry**

Cells (3-10x10^5 cells per stain) were washed with incubation buffer (2mM EDTA, 0.5% bovine serum albumin (BSA; Sigma-Aldrich) in PBS), spun down at 400 g for 5 mins at 4°C, and blocked in incubation buffer containing 10% rat serum (BioRad) and 10% mouse FcR blocking reagent (Miltenyi) on ice for 15 minutes. The antibodies were added directly to the cells or, when multiple antibodies with tandem fluorophores were included in the panel, the cells were washed with 1ml incubation buffer and spun down, before resuspending in the antibody mix containing 50µL Brilliant stain buffer (BD Horizon). After 40 minutes incubation on ice, the samples were washed with 1ml incubation buffer and resuspended in 200µL incubation buffer for analysis. Fluorescence minus one (FMOs) controls were used to set the gates. For intracellular staining of IL-12 the cells were fixed and permeabilized using intracellular (IC) fixation buffer and IC permeabilization buffer (eBioscience) following manufacturer's instructions. For intracellular staining of TCF-1, BD Pharmingen Transcription Factor Buffer Set (BD Biosciences) was used following the manufacturer's instructions. Cells were analyzed on Cytoflex LX (Beckman Coulter). Data were quantified using Cytexpert V2.4 software (Beckman Coulter). Details of the antibodies are provided in the Table S1.

**ELISA**

The plates were coated with 50µL per well of capture antibody (anti-IL-12p40 C15.6, Biolegend; 2 ng/mL in PBS) at 37°C for 2-3 hours or overnight at 4°C, then washed. All washes were performed with 0.1% Tween in PBS (PBS-T) three times at RT. The plates were blocked with 1% BSA in PBS for 90 minutes at RT followed by a wash. The samples (1:10 dilution in PBS) and standards (0.01 – 1 ng/mL IL-12) were transferred into the wells (50µL per well) in duplicates, incubated overnight at 4°C, and then washed prior to the incubation with the detection antibody (anti-IL-12p40 C17.8, Biolegend; 80 ng/mL in PBS with 1% BSA) for 90 minutes at RT. After a wash, 50µL of streptavidin hydrogen peroxidase (Sigma-Aldrich; 5 µg/mL in 1% BSA in PBS) per well was added and incubated for 30 minutes at RT in the dark, followed by a wash. 3,3′,5,5′-Tetramethylbenzidine substrate solution (ThermoFisher) was used to develop the signal and the reaction was stopped with 50µL $H_2SO_4$ per well. The optical density at 405/570 nm was measured in a plate reader (TECAN). IL-12 concentration in samples was derived from the standard curve and normalized to the tumor supernatant volume and tumor weight. The experimenter was blinded to the experimental groups.

**RNA isolation, preparation of sequencing libraries and mRNAseq of murine dendritic cells**

RNA isolation was performed in house using Arcturus PicoPure RNA isolation Kit (ThermoFisher Scientific) and DNA was removed with DNAse (Qiagen) following the manufacturers protocol. Library generation was performed by Novogen. Briefly, mRNA was purified using poly-T oligo-attached magnetic beads. After fragmentation, the first strand complementary DNA (cDNA) was synthesized using random hexamer primers, followed by the second strand cDNA synthesis. End repair, A-tailing, adaptor ligation, size selection, amplification and purification were performed. Libraries were quantified with qubit and the size distribution was detected by bioanalyzer. Quantified libraries were pooled. The clustering of the index-coded samples was performed according to the manufacturer's instructions. Following cluster generation, the library preparations were sequenced on Illumina NovaSeq 6000 and paired-end reads were generated.

**Murine dendritic cells mRNAseq data analysis**

Data analysis was performed by Novogen. FASTQ reads were quality checked and filtered using an in-house perl script, discarding reads with adaptor contamination, low quality (Q<5 constituting more than 50% of the read), and over 10% uncertain nucleotides (n>10%). Reads mapping to the reference genome GRCm38/mm10 were carried out and gene model annotation files downloaded from genome website directly. Index of the reference genome was built and paired-end clean reads aligned to the reference genome using Hisat2 v2.0.5 (34). Feature Counts v1.5.0-p3 (35) was used to count the reads numbers mapped to each gene, and then FPKM (Fragments Per Kilobase of transcript sequence per Millions base pairs sequenced) of each gene was calculated based on the length of the gene and reads count mapped to this gene. Differential expression analysis of two conditions/groups was performed using the DESeq2Rpackage (1.20.0). The resulting P-values were adjusted using the Benjamini and Hochberg's approach for controlling the false discovery rate. Genes with an adjusted P-value<=0.05 found by DESeq2 were assigned as differentially expressed.

Gene Ontology (GO) KEGG and Reactome pathways enrichment analyses of differentially expressed genes were implemented by the clusterProfiler R package, in which gene length bias was corrected. GO terms with corrected P value less than 0.05 were considered significantly enriched by differentially expressed genes.

**Correlation of cDC1-associated gene expression levels with patient survival**

A re-analysis of data published by Fisher et al. (11) was performed to determine a correlation between *XCR1-BATF3* gene expression levels and OS post-craniotomy in melanoma patients with BrM. Data were available upon registration and request through European Genome-phenome Archive (EGA) repository (https://ega-archive.org/datasets/EGAD00001005046). Supplementary Table 1 from Fischer et al. was used to extract the 88 RNA sample IDs from the 'Brain' Accession Site as well as the post-craniotomy survival data for individual patients. EGA RNA bam files were sorted by read name (samtools sort -n) and converted to paired fastq (samtools fastq -n -1 R1.fq -2 R2.fq) with samtools tool kit (36). All 88 RNA libraries were then mapped against the Ensembl GRCh38 reference genome (release 109) through STAR aligner (37) with default parameters. Genome fasta file was downloaded from https://ftp.ensembl.org/pub/release-109/fasta/homo_sapiens/dna/Homo_sapiens.GRCh38.dna.primary_assembly.fa.gz and gtf annotation from https://ftp.ensembl.org/pub/release-109/gtf/homo_sapiens/Homo_sapiens.GRCh38.109.gtf.gz. STAR-generated sorted BAM files were used as input for stringtie2 (38) with default parameters, except for triggering the -e option (only estimate the abundance of given reference transcripts), which in turn required the gtf annotation (-G Homo_sapiens.GRCh38.109.gtf). Normalized expression values per gene (TPMs) calculated by stringtie2 was then used in further downstream analyses.

A TPM expression-based z-Score sum of *XCR1* (ENSG00000173578) plus *BATF3* (ENSG00000123685) was calculated for each patient (N=88) in order to create a new 'High' and 'Low' expression status label to be assessed along with 'OS_Months' in the OS analysis. The bottom 40% of the z-Score sum were labelled as 'Low' expression (n=33) and the top

60% were labelled as ‘High’ expression (n=55). Bioconductor R packages ‘survival’ and ‘survminer’ were employed for OS analysis. All tools described in this paragraph were run under the R environment version 4.4. The survival curves were generated in Graph Pad Prism v7.

A re-analysis of data published by Vareslija et al. (39) was performed to determine a correlation between *XCR1-BATF3* gene expression levels and breast cancer BrM-specific survival. The data were accessed via the npriedig GitHub repository (https://github.com/npriedig/jnci_2018/blob/master/brainMetPairs.salmon.cts.txt). Unfiltered counts per million (CPM) for each sample were converted to log(CPM +1) and then scaled individually for *BATF3* and *XCR1* to produce *z*-scores. The sum of the data was taken to produce a dual *BATF3-XCR1* BrM gene signature for 21 samples (7M_RCS’ was omitted due to no available clinical data) and cross-referenced with survival post-brain metastasis (SPBM), as described in Table 1 of Varešlija et al. (39). The *BATF3-XCR1* dual gene signatures for each sample were ranked and split by quartile. Signatures in the upper quartile were classified as ‘high’ expression and signatures within the lower three quartiles were collectively classified as ‘low + intermediate’ expression. The Kaplan-Meier plot and statistical analysis was performed using GraphPad Prism version 10.2.1 (GraphPad Software, LLC).

**Reanalysis of scRNAseq data from melanoma brain metastases**

Raw scRNA-sequencing data published by Alvarez-Breckenridge et al. (19) were accessed through dbGaP. Alignment and expression quantification were performed using RSEM (v.1.3.3) and STAR (v.2.5.0a). Seurat (v.5.2.1) was used for subsequent filtering for high-quality single-cells included the exclusion of cells expressing <1000 genes and cells with > 20% of mitochondrial gene counts. Only cells expressing the *PTPRC* gene were included for downstream analysis. The NormalizeData function in Seurat was applied to normalise gene expression profiles using the LogNormalize method. The top 10,000 highly variable genes were identified with the FindVariableFeatures function. The data was scaled by utilising the

ScaleData function followed by principal component analysis on the scaled data with the RunPCA function, using default parameters. A shared nearest neighbour graph was constructed using the FindNeighbours function and cells were clustered using the Louvain algorithm with the FindClusters function. Finally, the RunUMAP function facilitated the visualization of all cells. The Panopticon signature scoring method was applied using signatures from Alavarez-Breckenridge et al. (19) to annotate individual cell types. Cells expressing at least one CD3 gene (*CD3D*, *CD3E* or *CD3G*) were included for downstream analysis. Cells with "CD4+ T-cell", "CD4+ T-helper cell" and "Treg" annotations were removed, followed by re-clustering of the data.

Effector-like, progenitor-exhausted and terminally-exhausted T-cell signatures derived from Miller et al. (10) were applied to the remaining CD8+ T cells using the AddModuleScore function in Seurat. For each cell, normalized scores were computed by dividing raw scores by the median across all cells for the respective signature. A dominant cell identity was assigned to each cell based on the highest normalised score. The code used in this study is available upon request.

**Statistical Analysis and Experimental Group Sizes**

Animal experimental group sizes were determined with power analysis (power = 80%, significance level = 5%, difference to be detected between groups = 50%) using the mean values and standard deviations (SDs) from previous our work (13). All animals were included in the survival analysis and tumor growth analysis. In cases where tumors were too small to obtain sufficient material for FACS analysis, the animals were excluded from this analysis. Statistical analysis was carried out using Graph Pad Prism v7. Data was plotted with mean and SD shown. All data was first assessed for statistical outliers using ROUT method with Q=0.1%, and the outliers excluded for further analysis. Data was then assessed for normal (Gaussian) distribution using Anderson Darling Test, significance level 0.05. Data was then analyzed using the most appropriate statistical method based on number of groups to be analyzed and normality of distribution. The number of data points, number of independent experiments for each graph and the statistical method used is stated in figure legends.

**Data sharing**

Murine cDC1 mRNAseq data generated in this study are available at the Gene Expression Omnibus (GEO) with accession number GSE308095.

### Supporting Figures

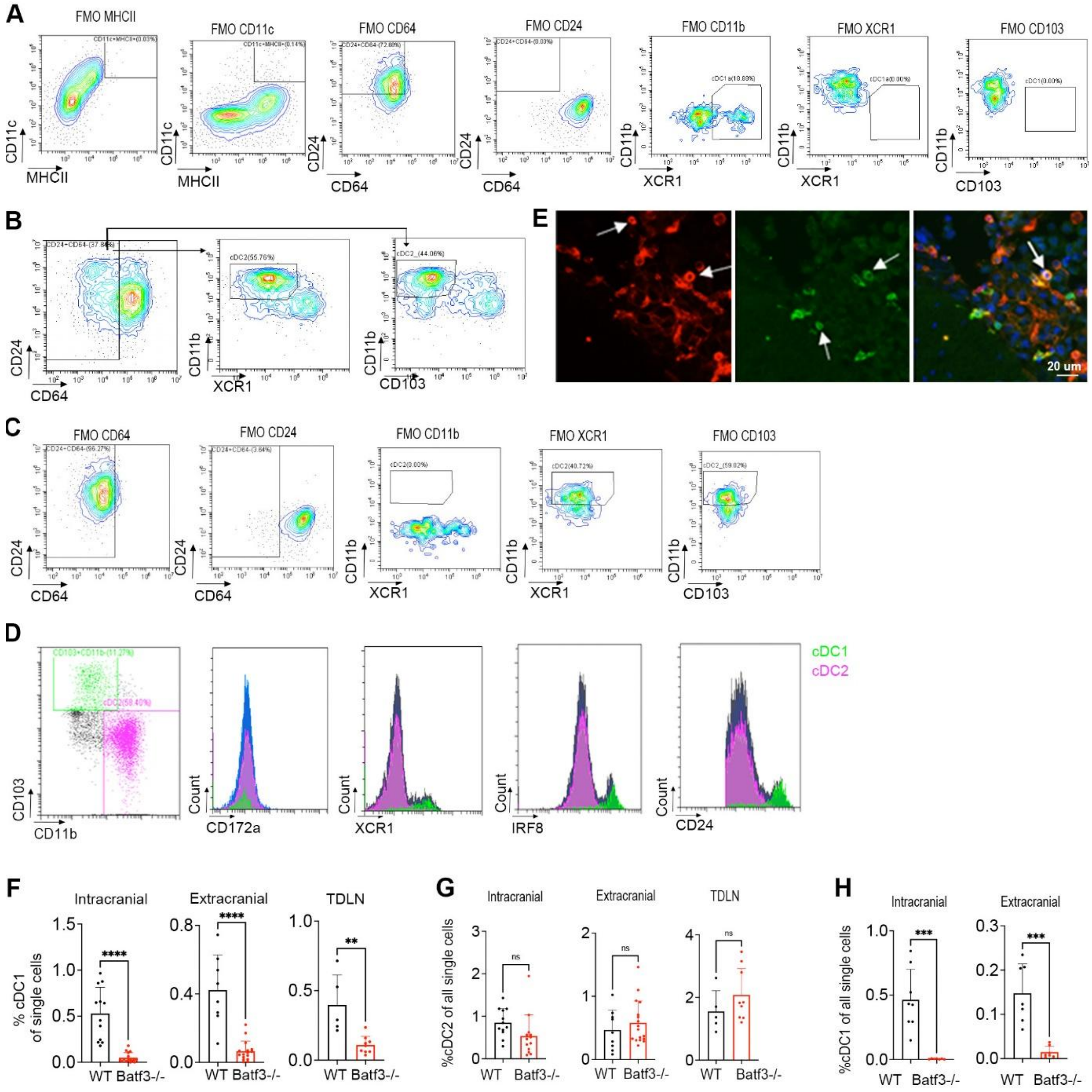


**Figure S1. Analysis of DCs by flow cytometry and immunofluorescence in the B16-OVA two-site model. (A)** Flow cytometry plots showing fluorescence minus one (FMOs) controls for the flow cytometry analysis in Fig. 1C. **(B)** Gating strategy for identification of cDC2 population by flow cytometry. CD45+ population was gated on MHCII+CD11c+ and then on CD64- population. Within the latter, the cDC2 population was defined as CD11b+CD103- or CD11b+XCR1-. **(C)** Flow cytometry plots showing FMOs controls for the flow cytometry analysis in B. **(D)** Representative flow cytometry plots of CD45+, MHCII+CD11c+, CD64-, CD24+Ly6C/Ly6G- cells infiltrating intracranial tumors, gated on cDC1 (green) and cDC2 (pink) populations, showing the expression of confirmatory markers; CD172a, XCR1, IRF8 and CD24. **(E)** Immunofluorescence staining of B16-OVA melanoma intracranial tumors isolated from the two-site model for MHCII (red) and CD103 (green) plus

DAPI nuclear stain (blue). **(F, G)** Quantification of cDC1s (F) and cDC2s (G) within single cells in B16-OVA tumors and TDLNs in WT and Batf3-/-mice. Intracranial tumors: n = 12 WT, n = 13 Batf3-/-; extracranial tumors (3 experiments pooled): n = 1/2/5 WT, n = 3/3/11 Batf3-/-; TDLNs (2 experiments pooled): n = 2/3 WT, n = 3/6 Batf3-/-. **(H)** Percentages of cDC1s of single cells in the PyMT two-site model (direct tumor implantation). Intracranial: n = 8 WT, n = 7 Batf3-/-; extracranial: n = 7 WT, n = 7 Batf3-/-. Significant differences in **F** - **H** were determined by two-tailed Student's T-test with unequal variance (* < 0.05; ** < 0.01; *** < 0.005; **** < 0.001).

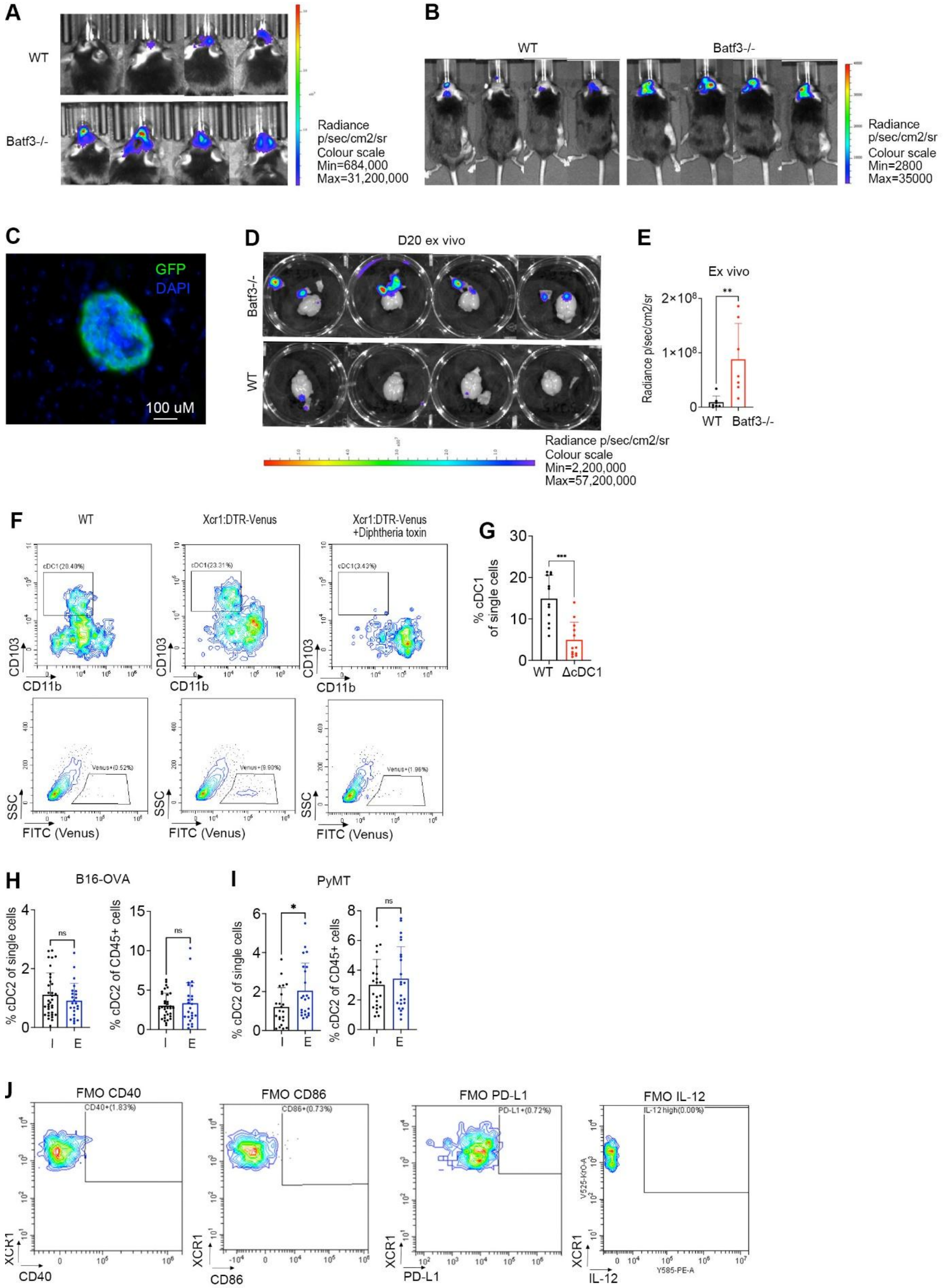


**Figure S2. Quantification of tumor growth and DCs. (A and B)** Representative bioluminescence images from the RET two-site model relating to Fig. 1G (A) and PyMT intracranial tumor implantation model relating to Fig. 1I (B). **(C)** An example of an intracranial GFP+ PyMT tumor lesion in the hematogenous PyMT two-site model. **(D)** Representative bioluminescence images from E. **(E)** Ex vivo quantification of whole-brain bioluminescence signal intensity in the hematogenous PyMT two-site model on day 20; n = 8 WT, n = 7 Batf3-/-. **(F)** Representative flow cytometry plots showing DT-induced depletion of the cDC1 population in Xcr1:DTR-Venus mice compared to WT controls, as detected by gating on the CD11b-CD103+ cells within CD45+CD11c+MHCII+CD24+CD64- population (**top**) or on the Venus+ population within the single cell gate (**bottom**). **(G)** Percentages of cDC1s within single cells as quantified by flow cytometry within intracranial B16-OVA melanoma tumors in WT and Xcr1:DTR-Venus (ΔcDC1) mice. Data from single experiment; n = 12 per group. **(H and I)** Percentages of cDC2s within single cells or within CD45+ cells as quantified by flow cytometry within intracranial (I) and extracranial (E) tumors in the B16-OVA two-site model (4 pooled experiments; intracranial: n = 7/7/6/6; extracranial: n = 7/4/7/1) (H) and in the PyMT two-site model with direct intracranial tumor implantation (2 pooled experiments; intracranial: n = 8/6; extracranial: n = 6/9) (I). **(J)** Flow cytometry plots showing fluorescence minus one (FMOs) controls for the flow cytometry analysis in Fig. 2D. Significant differences in **E** and **G** were determined by two-tailed Student's T-test with unequal variance, and significant differences in **H** and **I** by Mann Whitney Test (* < 0.05; ** < 0.01; *** < 0.005; **** < 0.001).

**A**

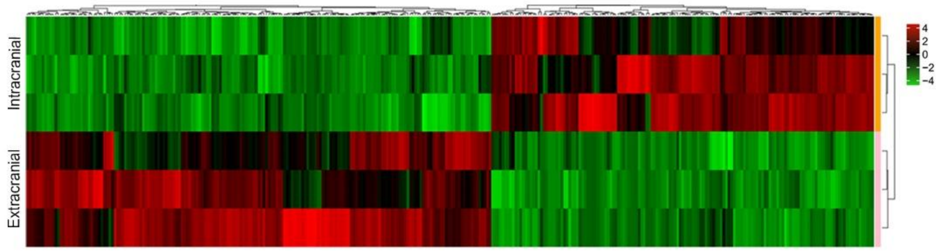


**B**

| Category | GOID | Description | GeneRatio | BgRatio | pvalue | padj | geneName | Count |
|---|---|---|---|---|---|---|---|---|
| BP | GO:0070371 | ERK1 and ERK2 cascade | 40/921 | 296/16289 | 2.67e-07 | 8.04e-05 | Ccl6/Ccl7/P2ry1/Myc/Kdr/Dstyk/Fgfr1/Ephb4/App/Dusp3/Ptpn1/Ccl8/Dnajc27/Garem1/Mif/Erbb2/Ddr1/Ccl9/Ctsh/Tlr4/Agt/Atp6ap1/Marco/Pdgfc/Cd44/Rps6ka6/Epha3/Cyr61/Timp3/C5ar1/Ccl5/Gcg/Ctgf/Pla2g5/Ryk/Hras/Fgf21/Xcl1/Dab2 | 40 |
| BP | GO:0043410 | positive regulation of MAPK cascade | 53/921 | 450/16289 | 3.13e-07 | 8.86e-05 | Ccl6/Ccl7/P2ry1/Myd88/Kdr/Dstyk/Fgfr1/Ephb4/App/Ptpn1/Ccl8/Map3k6/Tgfb3/Dnajc27/Garem1/Gadd45g/Mif/Lpar3/Erbb2/Itga1/Ddr1/Ccl9/Gadd45a/Tlr4/Avpi1/Stk25/Agt/Atp6ap1/Marco/Wwc1/Pdgfc/Cd44/Epha3/C5ar1/Ccl5/Gcg/Plcb1/Ctgf/Tdgf1/Traf2/Pla2g5/Ryk/Tpd52l1/Tnfrsf10b/Hras/Fgf21/Xcl1/Fzd4/Dab2/Fcgr2b/Map3k3/Zeb2/Ddt | 53 |
| BP | GO:0070374 | positive regulation of ERK1 and ERK2 cascade | 31/921 | 210/16289 | 8.77e-07 | 0.000183412 | Ccl6/Ccl7/P2ry1/Kdr/Dstyk/Fgfr1/Ephb4/App/Ccl8/Dnajc27/Garem1/Mif/Erbb2/Ddr1/Ccl9/Tlr4/Atp6ap1/Marco/Pdgfc/Cd44/Epha3/C5ar1/Ccl5/Gcg/Ctgf/Pla2g5/Ryk/Hras/Fgf21/ | 31 |
| BP | GO:0070372 | regulation of ERK1 and ERK2 cascade | 37/921 | 281/16289 | 1.41e-06 | 0.000220815 | Ccl6/Ccl7/P2ry1/Kdr/Dstyk/Fgfr1/Ephb4/App/Dusp3/Ptpn1/Ccl8/Dnajc27/Garem1/Mif/Erbb2/Ddr1/Ccl9/Tlr4/Atp6ap1/Marco/Pdgfc/Cd44/Rps6ka6/Epha3/Cyr61/Timp3/C5ar1/Ccl5/Gcg/Ctgf/Pla2g5/Ryk/Hras/Fgf21/Xcl1/Dab2/Ddt | 37 |
| BP | GO:0002755 | MyD88-dependent toll-like receptor signaling pathway | 8/921 | 20/16289 | 6.93e-06 | 0.00068022 | Cd300a/Myd88/Tlr1/Tlr8/Tlr4/Irak2/Tlr5/Tlr7 | 8 |

**C**

| Category | GOID | Description | GeneRatio | BgRatio | pvalue | padj | geneName | Count |
|---|---|---|---|---|---|---|---|---|
| BP | GO:0035455 | response to interferon-alpha | 10/834 | 23/16289 | 7.30E-08 | 0.000169049 | Ifi204/Ifit2/Axl/Adar/Bst2/Eif2ak2/Ifnar1/Klhl20/Gas6/Ifnar2 | 10 |
| BP | GO:0032647 | regulation of interferon-alpha production | 8/834 | 27/16289 | 4.24E-05 | 0.021837342 | Irf7/Havcr2/Ifih1/Tlr3/Chuk/Setd2/Stat1/Zc3hav1 | 8 |
| BP | GO:0032607 | interferon-alpha production | 8/834 | 28/16289 | 5.68E-05 | 0.02628785 | Irf7/Havcr2/Ifih1/Tlr3/Chuk/Setd2/Stat1/Zc3hav1 | 8 |
| BP | GO:0032727 | positive regulation of interferon-alpha production | 7/834 | 22/16289 | 7.79E-05 | 0.028174626 | Irf7/Ifih1/Tlr3/Chuk/Setd2/Stat1/Zc3hav1 | 7 |
| BP | GO:0034340 | response to type I interferon | 9/834 | 37/16289 | 7.88E-05 | 0.028174626 | Irf7/Isg15/Adar/Ifnar1/Nlrc5/Stat2/Setd2/Ifnar2/Stat1 | 9 |
| BP | GO:0035457 | cellular response to interferon-alpha | 5/834 | 11/16289 | 1.24E-04 | 0.028268872 | Ifi204/Ifit2/Axl/Ifnar1/Gas6 | 5 |

**D**

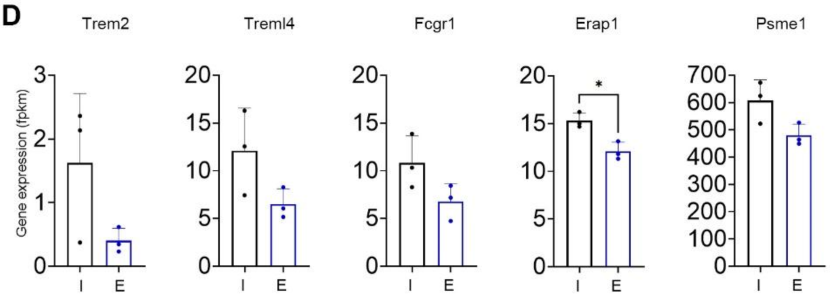


**Figure S3. Analysis of cDC1 transcriptional profiles (A)** A heat map displaying differentially expressed genes between cDC1s isolated from intracranial and extracranial tumors in the two-site B16-OVA model (FDR≤0.05). **(B)** gene set enrichment analysis (GSEA) using GO database revealed a significant downregulation of pathways related to the positive regulation of ERK1 and ERK2, positive regulation of MAPK, and MyD88-dependent toll-like receptor signaling pathway in intracranial cDC1s. **(C)** GSEA using GO database revealed a significant upregulation of pathways related to type-I-interferon / IFNα in intracranial cDC1s. Significant differences in D were determined by two-tailed Student's T-test with unequal variance (* < 0.05; ** < 0.01; *** < 0.005; **** < 0.001).

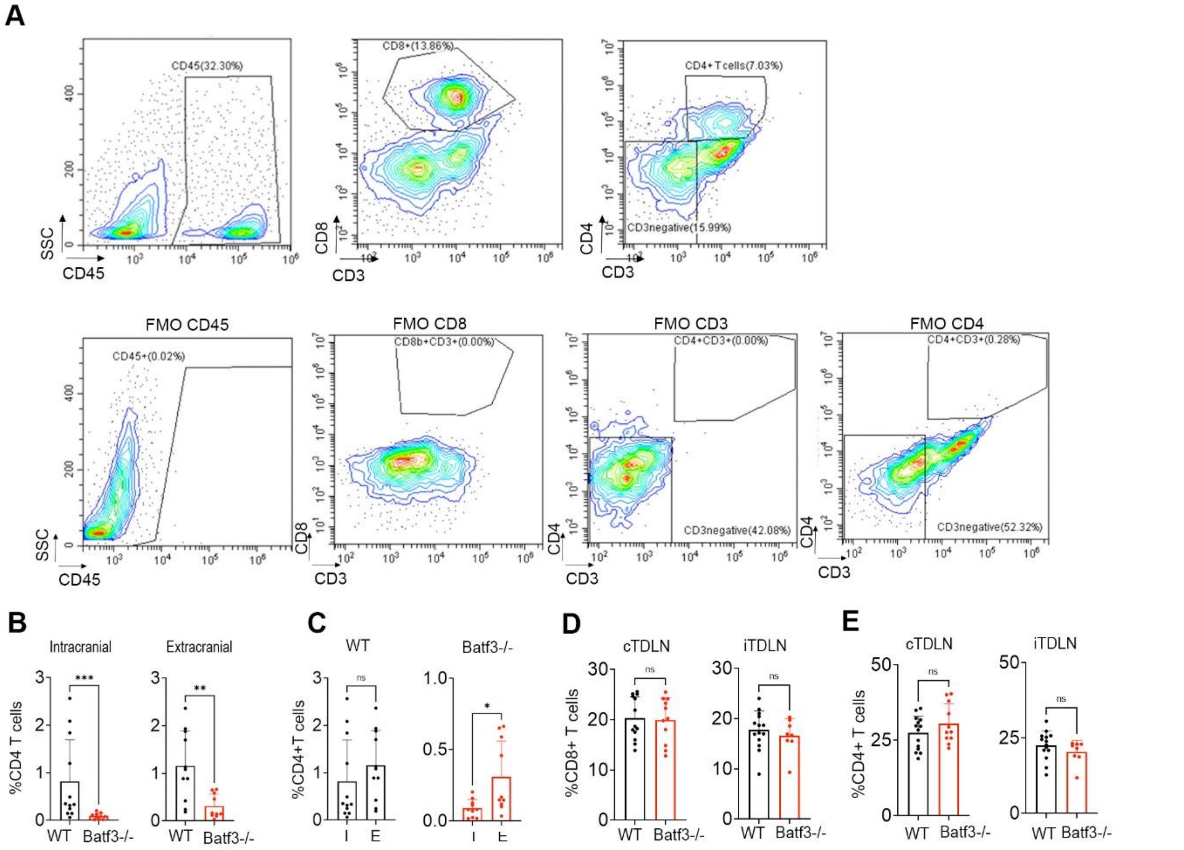


**Figure S4. Analysis of T cells by flow cytometry. (A)** Representative flow cytometry plots showing the gating strategy for CD4+ and CD8+ T cells (top row) and corresponding FMO controls (bottom row). **(B)** Percentages of intra-tumoral CD4+ T cells within single cells in the two-site B16-OVA model (2 pooled experiments; intracranial; n = 5/7 WT, n = 3/7 Batf3-/-; extracranial: n = 5/6 WT, n = 4/6 Batf3-/-. **(C)** data from B were replotted for direct comparison of percentages of CD4+ T cells between the intracranial (I) and extracranial (E) tumors. **(D, E)** Percentages of CD8+ (D) and CD4+ T cells (E) within single cells in cTDLNs and iTDLNs in the two-site B16-OVA model (2 pooled experiments; cTDLNs: n = 8/6 WT, n = 8/3 Batf3-/-; iTDLNs: n = 8/6 WT, n = 6/2 Batf3-/-). Significant differences **B** and **C** were determined by Mann Whitney Test, and in **D** and **E** by two-tailed Student's T-test with unequal variance (* < 0.05; ** < 0.01; *** < 0.005; **** < 0.001).

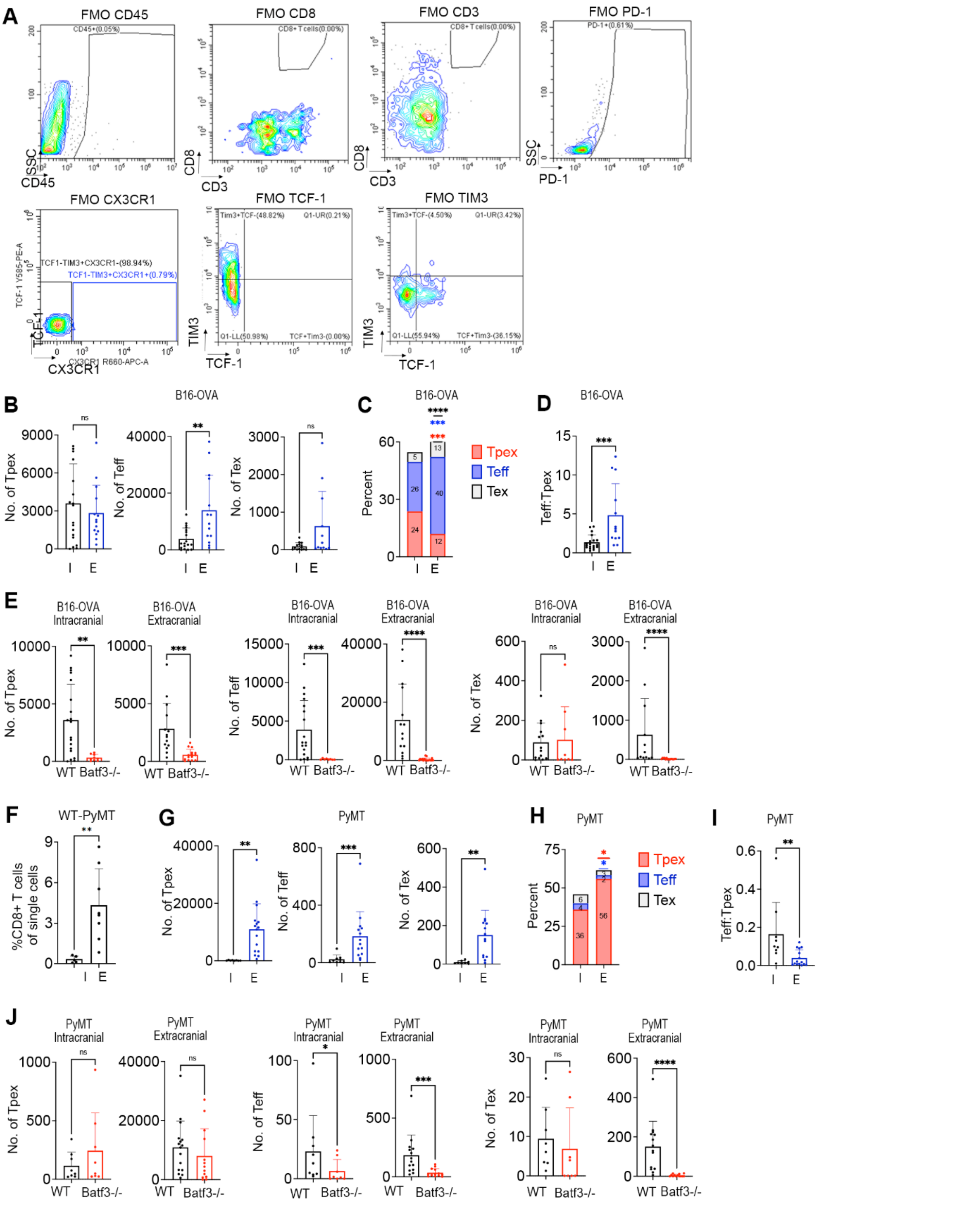


**Figure S5. Analysis of exhausted CD8+ T cell subsets by flow cytometry. (A)** Flow cytometry plots showing fluorescence minus one (FMOs) controls for the flow cytometry analysis in Fig. 4H. **(B-D)** Replotted data from (E) for direct comparison of the Tpex, Teff and Tex numbers per 100,000 single cells (B), their proportions within CD8+ T cells (C), and Teff/Tpex ratios (D) between intracranial (I) and extracranial (E) tumors in the two-site B16-OVA model in the WT mice. **(E)** Numbers of intra-tumoral Tpex, Teff and Tex per 100,000 single cells in the two-site B16-OVA model, as quantified by flow cytometry. **(F)** Percentages of CD8+ T cells within single cells in intracranial (I) and extracranial (E) tumors in the two-site PyMT model with direct intracranial tumor implantation in the WT mice (n = 6 intracranial, n = 8 extracranial). **(G-I)** Comparison of the Tpex, Teff and Tex numbers per 100,000 single cells (G), their proportions within CD8+PD1+ T cells (H), and Teff/Tpex ratios (I) between intracranial (I) and extracranial (E) tumors in the two-site PyMT model with direct intracranial tumor implantation in the WT mice. **(J)** Numbers of Tpex, Teff and Tex per 100,000 single cells in intracranial and extracranial tumors in the two-site PyMT model in WT and Batf3-/- mice. Data in **G-J** were pooled from 2 independent experiments; intracranial: n = 5/5 WT, n = 4/4; Batf3-/-; extracranial: n = 8/8 WT, n = 8/7 Batf3-/-. Significant differences **B (Teff, Tex), C (Tex), D, E, I,** and **J** were determined by Mann Whitney Test. Significant differences in **B (Tpex), C (Teff and Tpex), F, G,** and **H** were determined by two-tailed Student's T-test with unequal variance (* < 0.05; ** < 0.01; *** < 0.005; **** < 0.001).

**Table S1.** Details of antibodies used in the study.

| **Antibody** | **Company, Catalog number, Clone** |
|---|---|
| CD45-BUV395 | (BD Biosciences Cat# 564279, RRID:AB_2651134) |
| CD11c-PerCPCy5.5 | (Thermo Fisher Scientific Cat# 45-0114-80, RRID:AB_925728) |
| MHCII-Vioblue | (Miltenyi Biotec Cat# 130-102-145, RRID:AB_2660060) |
| CD24-APC | (Miltenyi Biotec Cat# 130-103-372, RRID:AB_2656576) |
| CD11b-BV605 | (BioLegend Cat# 101237, RRID:AB_11126744) |
| CD64 (X54-5/7.1)-FITC | (BioLegend Cat# 139315, RRID:AB_2566555) |
| CD103 (M290)-PE | (BD Biosciences Cat# 561043, RRID:AB_10565963) |
| XCR1 (ZET)-APC/Cy7 | (BioLegend Cat# 148223, RRID:AB_2783117) |
| IRF8 (V3GYWCH)-PE/Cy7 | (Thermo Fisher Scientific Cat# 25-9852-82, RRID:AB_2784675) |
| CD45-BV605 | (BioLegend Cat# 103140, RRID:AB_2562342) |
| CD8b-PECy7 | (BioLegend Cat# 126616, RRID:AB_2562777) |
| CD3e-APC-Vio770 | (Miltenyi Biotec Cat# 130-121-440, RRID:AB_2801817) |
| NK1.1-FITC | (BioLegend Cat# 108706, RRID:AB_313393) |
| DX5(CD49b)-PE | (Miltenyi Biotec Cat# 130-123-702, RRID:AB_2811545) |
| CD11b-V450 | (BD Biosciences Cat# 560455, RRID:AB_1645266) |
| CD4-APC | (Miltenyi Biotec Cat# 130-116-546, RRID:AB_2727604) |
| CD172a (SIRP alpha)- BV510 | (BD Biosciences Cat# 740159, RRID:AB_2739912) |
| Ly-6G and Ly-6C -Alexa Fluor-700 | (BD Biosciences Cat# 557979, RRID:AB_396971) |
| CD64-PE/Cyanine7 | (BioLegend Cat# 139313, RRID:AB_2563903) |
| CD103-BV510 | (BD Biosciences Cat# 563087, RRID:AB_2721775) |
| CD274 (PD-L1)- Brilliant Violet 785 | (BioLegend Cat# 124331, RRID:AB_2629659) |
| CD40-PE-Cyanine5 | (Thermo Fisher Scientific Cat# 15-0401-82, RRID:AB_468747) |
| CD86-APC-eFluor 780 | (Thermo Fisher Scientific Cat# 47-0862-82, RRID:AB_2815162) |
| CD80-BUV661 | (BD Biosciences Cat# 741515, RRID:AB_2870964) |
| IL-12 (p40/p70)-PE | (BD Biosciences Cat# 554479, RRID:AB_395420) |
| XCR1-BV510 | (BioLegend Cat# 148218, RRID:AB_2565231) |
| TCF-7/TCF-PE | (BD Biosciences Cat# 564217, RRID:AB_2687845) |
| CD366 (TIM3)-FITC | (Thermo Fisher Scientific Cat# 11-5870-82, RRID:AB_2688129) |
| CX3CR1-APC | (BioLegend Cat# 149008, RRID:AB_2564492) |
| PD-1-BV421 | (BioLegend Cat# 135221, RRID:AB_2562568) |
| Anti-CD103 | (R and D Systems Cat# AF1990, RRID:AB_2128618) |
| anti-MHCII | (BD Biosciences Cat# 556999, RRID:AB_396545) |
| anti-GFP | (Abcam Cat# ab13970, RRID:AB_300798) |
| anti-chicken-FITC | (Jackson ImmunoResearch Labs Cat# 703-096-155, RRID:AB_2340357) |
| anti-Goat IgG (H+L) Alexa Fluor Plus 647 | (Thermo Fisher Scientific Cat# A32849, RRID:AB_2762840) |
| anti-rat-Cy3 | (Millipore Cat# AP189C, RRID:AB_92645) |
